# An in vivo CRISPR screen in chick embryos reveals a role for MLLT3 in specification of neural cells from the caudal epiblast

**DOI:** 10.1101/2024.05.16.594506

**Authors:** Ashley RG Libby, Tiago Rito, Arthur Radley, James Briscoe

## Abstract

Tissue development relies on the coordinated differentiation of stem cells in dynamically changing environments. The formation of the vertebrate neural tube from stem cells in the caudal lateral epiblast (CLE) is a well characterized example. Despite an understanding of the signalling pathways involved, the gene regulatory mechanisms remain poorly defined. To address this, we developed a multiplexed in vivo CRISPR screening approach in chick embryos targeting genes expressed in the caudal epiblast and neural tube. This revealed a role for *MLLT3*, a component of the super elongation complex, in the specification of neural fate. Perturbation of *MLLT3* disrupted neural tube morphology and reduced neural fate acquisition. Mutant forms of Retinoic Acid Receptor A lacking the *MLLT3* binding domain similarly reduced neural fate acquisition. Together, these findings validate an in vivo CRISPR screen strategy in chick embryos and identify a previously unreported role for *MLLT3* in caudal neural tissue specification.

## INTRODUCTION

To generate functioning tissues during development, a wide variety of cell types must be specified in a timely and organized manner. As such, individual cells transition through multiple embryonic regions (*1–3*), experience varying signalling regimes (*4–8*), and activate distinct gene expression programmes as they differentiate towards their eventual fate (*9–11*). Across tissues, developmental stages, and species, coordinating signalling processes with gene expression is necessary to ensure the robust and precise outcome of tissue development (*12–14*).

The progressive elaboration of the vertebrate trunk from the caudal lateral epiblast (CLE) exemplifies a dynamic signalling landscape coordinating the acquisition of diverse cell fates. Here, a multipotent population of cells, termed neuromesodermal progenitors (NMPs), gives rise to successively more posterior neural and paraxial mesodermal (PSM) lineages (*15–18*) while proliferation of NMPs fuels the elongation of axial tissues (*17, 19, 20*). Consequently, the rates at which NMPs emerge, self-renew, and differentiate must be closely regulated within the CLE to balance the production of trunk tissues. To this end, the interplay between WNT, FGF, and Retinoic Acid (RA) signalling regulates the behaviors of NMPs and other caudal stem cell populations (*4, 15, 21, 22*). Perturbations or spatial redistribution of these signals can delay or alter the acquisition of subsequent cell fates. For example, protracted FGF signalling impedes neural differentiation (*23–25*), RA accelerates neural fate acquisition (*26, 27*), and elevated WNT signalling drives mesodermal fates (*28, 29*). Despite an overall understanding of the signalling molecules involved in balancing the allocation of NMP-derived trunk fates (*30, 31*), our current knowledge of the specific gene regulatory mechanisms involved in this fate decision is based on a handful of genes, leaving a gap in knowledge of novel mechanisms and genes governing the rapid transitions involved in fate decisions.

In recent years, pooled CRISPR/Cas9 genetic screens targeting multiple genes have proved valuable tools for the interrogation of gene regulatory mechanisms (*32–36*). However, only a limited number of *in vivo* pooled screens have been performed to date, in part due to technical challenges such as the even distribution of guides, low transfection rate *in vivo*, and low cell numbers due to tissue size. As such, *in vivo* pooled screens have been restricted to more easily accessible tissues such as the brain (*37–41*), or to species such as zebrafish that are more experimentally tractable (*42*). Nevertheless, the outcomes of *in vivo* screens have demonstrated their potential to identify previously understudied genes involved in specific processes (*43, 44*). Given the potential of in vivo screens, there is a need for the development of methods that allow targeting of specific tissues across multiple species.

To refine our knowledge of the molecular mechanisms that govern neural tube development, we developed a pooled *in vivo* CRISPR screen in chick embryos. Genetic targets were identified using a single-cell transcriptome dataset (*45*) and perturbed following electroporation of CRISPR guides into the chick CLE in a pooled fashion. Post perturbation, enrichment or depletion of lineages arising from the CLE were assayed by single-cell RNA sequencing. Our results emphasized a multifaceted regulation of the differentiation of CLE cells to neural tube fate that involves the interaction of FGF, WNT, and RA signalling with gene regulatory programmes. Further, we identified a previously unobserved role of the gene *MLLT3*, a member of the super elongation complex, as a key factor in controlling the behaviour of cells in the CLE. While observed *MLLT3* expression was restricted to the CLE and primitive streak, targeting *MLLT3* resulted in a reduction of knockout (KO) cells both within the CLE and within the neural tube. We provide evidence that this reduction in neural fate was due to an interaction with Retinoic Acid Receptor A (RARα). Overall, this work demonstrates a strategy for pooled perturbation screens in chick embryos, highlighting the utility of *in vivo* pooled screens to generate a mechanistic understanding of gene regulation, and provides new insight into how RA signalling is coupled with spatially defined gene expression to facilitate rapid fate acquisition and ensure the proportioned generation of tissues.

## RESULTS

### Identification of genes marking the transition from caudal lateral epiblast to neural fate

To investigate the control of fate transitions from the CLE to neural tube, we analyzed single-cell transcriptome datasets of the chick embryo at stages HH10 and HH11 (*45*) (Figure 1A). We computationally isolated cells annotated as primitive streak, CLE, pre-neural tube, and neural tube based on their gene expression (Figure 1B) and used an Entropy Sorting derived feature selection algorithm (cESFW) (*46, 47*) to identify co-regulated genes (total of 2505) in the trajectory from CLE to neural tube (Figure S1A,B, Sup. Table 1). These genes were then used to subset the scRNA-seq data (Figure 1B - right) and identified twelve clusters of cells (Figure 1B,D) in which the pattern of gene expression was associated with anatomical regions of the embryo (Figure 1C, S1D), including a population of *SOX2, SOX3,* and *TBXT* co-expressing cells representing presumptive NMPs (Figure S1D lower right). A pseudotime algorithm generated predicted lineage trajectories (Figure 1E, S1C), providing higher resolution of cell states and highlighting several previously under-studied genes. For example, expressed of *MLLT3* overlapped with *TBXT and MSX1* within the primitive streak and CLE; and *F2RL1* and *CMTM8* were expressed in the CLE with *CLDN1* and *NKX1-2* (Figure 1E).

**Figure 1:**
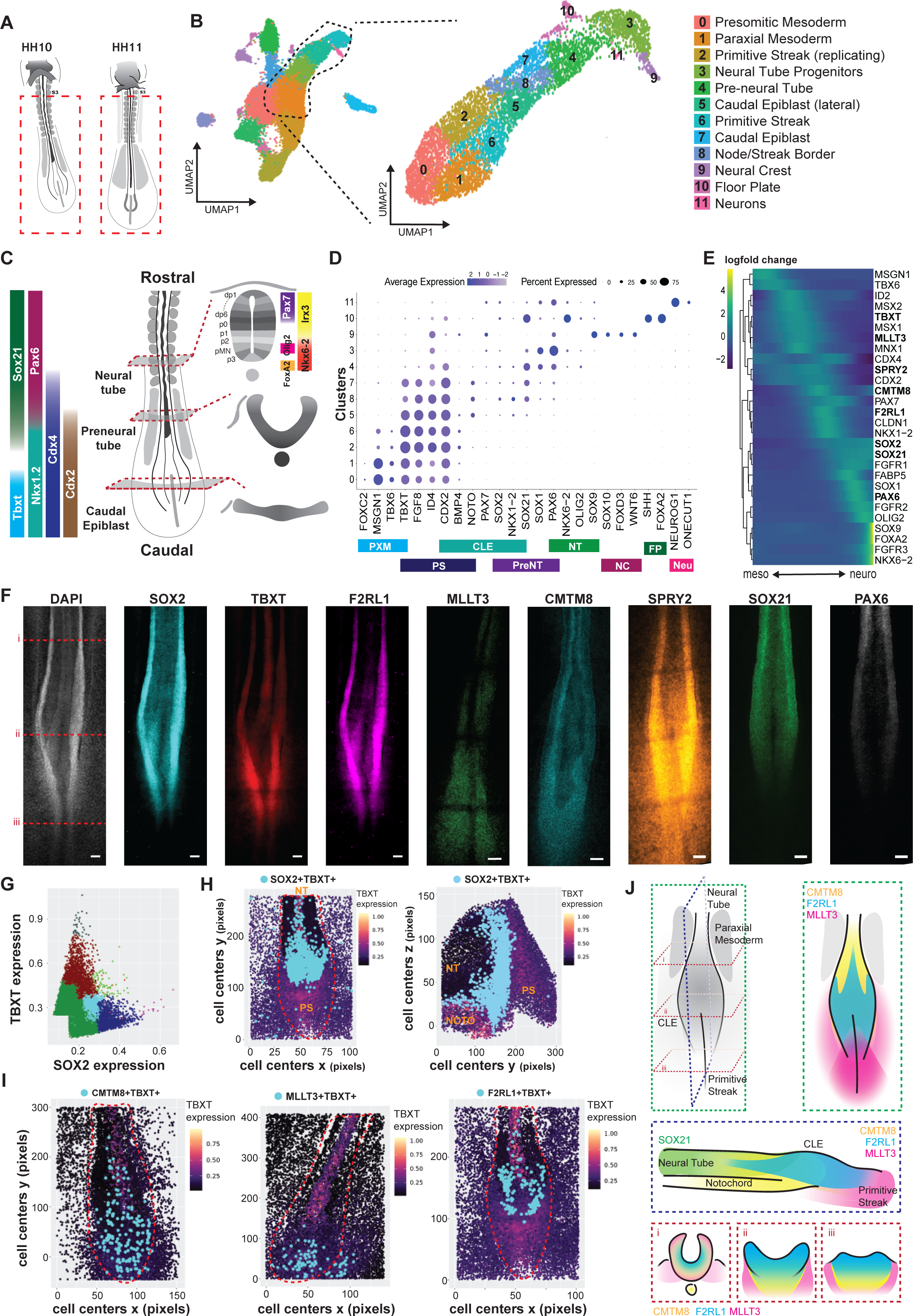
Mapping gene expression in CLE and neural tube of chick embryos. A) Schematic of embryos used in previously published dataset (*102*). B) Computational selection and re-clustering of dataset focusing on the primitive streak, caudal lateral epiblast (CLE), and neural tube. C) Adapted schematic (*103*) indicating known transcription factor expression in the primitive streak, CLE and neural tube. D) Dot plot of representative gene expression across dataset of lineage markers. PXM – paraxial mesoderm, PS – primitive streak, CLE – caudal lateral epiblast, PreNT – preneural tube, NT – neural tube, NC – neural crest, FP – floor plate, Neu – mature neurons. E) Heatmap of pseudo-trajectory from mesoderm to neural. Gene expression in log(10) counts. F) Example HCRs in HH10 chick embryos of indicated genes in Figure 1E (i indicates pre neural tube, ii indicated CLE, iii indicates primitive streak). G) Hierarchical clustering of TBXT transcript count with SOX2, CMTM8, or F2RL1 transcript count (light blue dots marks population plotted in Figure 1H. H) *LEFT:* Population of TBXT+SOX2+ cells (lightblue) that are identified within the black circle from Figure 1G were mapped onto *TBXT* expression maps of the imaged embryo in the xy plane where rostral is top and caudal is bottom (red outline demarks primitive streak – PS, CLE, and neural tube – NT). *RIGHT:* Population of TBXT+SOX2+ cells (lightblue) that are identified within the black circle from Figure 1G mapped onto *TBXT* expression maps of the imaged embryo in the yz plane where rostral is left and caudal is right. (primitive streak – PS, notochord - NOTO, and neural tube – NT. I) Populations of CMTM8+ TBXT+, MLLT3+TBXT+, and F2RL1+TBXT+ cells (lightblue) mapped to TBXT expression maps of HH10 embryos where the top is rostral and bottom is caudal (red outline demarks the primitive streak, CLE, and neural tube). J) Schematic of expression domains determined via HCR. Green box mark view from above (xy). Blue box marks view from side (yz). Red boxes mark three cross-sectional (xz) views from rostral to caudal. (Scale bar, 50um.)

To refine the map of *CMTM8*, *MLLT3*, and *F2RL1* expression, we examined expression of cytoskeletal components reported to mark various domains of the CLE (*45*): *CNTN2* (preneural), *NEFM* (node-streak border), and *ADAMTS18* (primitive streak) in the wildtype dataset (Figure S1F). *CMTM8* largely overlapped with *NEFM* in the CLE, although the expression domain of *CMTM8* extends beyond that of the *NEFM* domain. *MLLT3* overlapped with ADAMTS18 in the primitive streak and with NEFM in the CLE, again representing a larger domain than the cytoskeletal component. Finally, *F2RL1*, had an expression domain similar to *NEFM*, largely restricted to the CLE.

To validate the observed gene overlap and spatial location of the transcriptome analysis, we performed hybridization chain reaction (HCR) in HH10 embryos (Figure 1F). We quantified gene expression (Figure S2) and used hierarchical clustering of cell expression profiles to identify cell populations within the CLE. Co-expression of *SOX2* and *TBXT* was identified in a population of putative NMPs within the CLE (Figure 1G – light blue dots). These could be mapped at single cell resolution back to the embryo (Figure 1H – light blue dots in xy axis *(left)* and yz axis *(right)*). To define the spatial-transcriptomic arrangement of the CLE, the expression profiles of *CMTM8*, *F2RL1*, *MLLT3*, and *SOX21* (a non-CLE expressed control) were compared to that of *TBXT* (Figure S3A) and mapped to the embryo (Figure 1I, S3B-D). This produced a refined CLE domain map (Figure 1J) of *CMTM8*, *F2RL1*, and *MLLT3* expression in the CLE where each have spatially distinct cell populations co-expressing *TBXT* (Figure 1I, S3C-E). *CMTM8+TBXT+* cells were in the CLE with a minor contribution to the notochord (Figure 1I: left - blue dots, S3C: right - blue dots). *MLLT3+TBXT+* cells were located solely in the CLE and primitive streak (Figure 1I: center - blue dots, S3D: left - blue dots). Cells with high levels of *F2RL1* were located sporadically outside *TBXT* regions (Figure S2C), while medium *F2RL1* expression largely overlapped with *TBXT* in the CLE (Figure 1I: right - blue dots, S3D: left - blue dots), and all three genes *(CMTM8*, *F2RL1*, and *MLLT3)* were lowly expressed in the neural tube (Figure S2C-E). Overall, this dataset highlights domain-specific patterns of gene expression in the CLE and neural tube that agree with previously reported datasets. These observations raise the question of whether these genes regulate morphogenic signal interpretation to control balanced lineage emergence.

### Adapting chick CRISPR/CAS9 system for pooled single-cell RNA sequencing screening

To investigate the function of these identified transition-associated genes expressed in the CLE, we conducted an *in vivo* pooled CRISPR screen in chick embryos. To this end, we modified a previously published, chicken-specific CRISPR system (*48, 49*). The system utilises dual plasmids; one plasmid encoded a CAG-driven Cas9 and a Citrine reporter and the second contained the chick U6.3 promoter driving a guide RNA (gRNA) in which we inserted a capture sequence in the stem loop, to allow detection in 10X single-cell RNA-seq (*50*)(Figure 2A).

**Figure 2:**
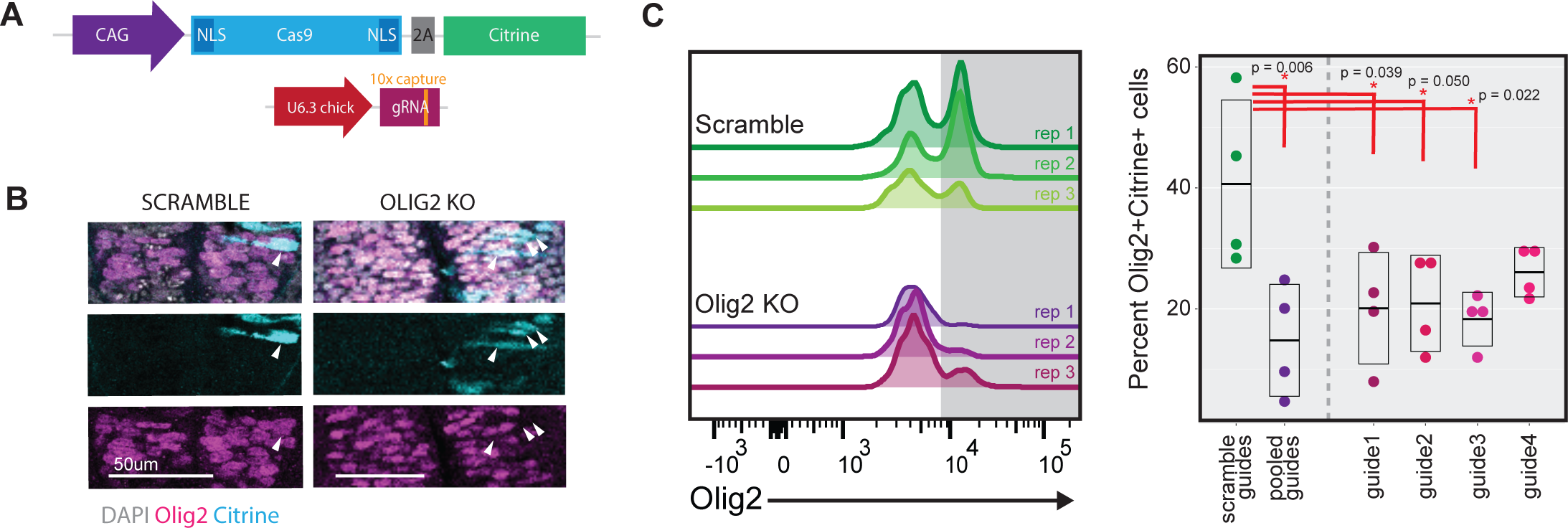
Chick in ovo CRISPR system. A) Schematic of the two-plasmid CRISPR system. CAG driven CAS9-IRES-Citrine and U6 driven gRNA with additional capture sequence for 10X feature barcoding capture. B) Immunofluorescent images for the indicated makers in transverse sections of the neural tube 24h post electroporation. White arrows mark detection of Cas9 and in KO condition loss of Olig2 protein. C) Flow cytometry analysis of Olig2 in cells from dissociated embryos 24h post neural tube electroporation (* represents p < 0.05 by ANOVA and subsequent Tukey tests, n = 4 embryos per condition).

To assess whether the capture sequence affected system performance, we designed four gRNAs targeting the transcription factor *OLIG2*, the knockout of which in the neural tube has been well documented (*51, 52*). The gRNAs and the Cas9 encoding plasmid were introduced into HH10 chick neural tubes via neural tube injection and unilateral electroporation. The side of the neural tube that received gRNAs (right) displayed disrupted Olig2 domains compared to the non-electroporated side (Figure S4A). Further, cells that had received the Cas9 plasmid lacked Olig2 protein (Figure 2B, S4A). By contrast, Olig2 expression remained unperturbed in control embryos electroporated with scramble gRNAs. To quantify the efficiency, we used flow cytometry to isolate Sox2+ neural progenitors from the neural tubes of embryos electroporated with either the scramble gRNA or Olig2 targeting gRNA pool (Figure S4B). This confirmed reduced Olig2 protein in electroporated cells from embryos receiving the Olig2 gRNA pool compared to the scramble control (Figure 2C left), suggesting that the capture sequence did not influence system performance.

With the aim of performing a pooled screen targeting multiple genes, we evaluated the efficacy of individual gRNAs in our perturbation system, as gRNA failure—failure to mutate the target gene—has been reported (*53, 54*). Flow cytometry of Sox2+ progenitors electroporated with individual Olig2 gRNAs revealed significantly reduced Olig2 levels from the majority of guides, however one gRNA (guide 4) did not deplete Olig2+ cells significantly (Figure 2C, right). Overall, we conclude that the changes made to the system, including the addition of the capture sequence to gRNAs, did not impair Cas9 activity. We proceeded with designing a screen in which unique barcodes paired with each guide variant allow comparison between guides as an internal quality control for guide failure.

### Generating an in vivo pooled screen for regulators of neural differentiation

To identify factors involved in the CLE-to-neural transition, we generated a pooled guide RNA (gRNA) library targeting genes selected from the transition-associated gene list. In total, the library consisted of 100 gRNAs targeting 25 candidate genes (4 gRNAs/gene) and two additional scramble controls (Table 2). This list included signaling receptors of pathway components from key pathways at play (*DUSP6, FGFR1, FGFR2, FGFR3, MAPK3 SPRY2, RARA, RARB, RARG, YAP1*), several reference genes (*TBXT*, *PAX7*, *OLIG2)*, the knockout of which have well studied phenotypes (*51, 52, 55–59*), and CLE transition genes identified by cESFW (*CLDN1, CMTM8, EMX2, F2RL1, FABP5, GREB1, KRAS, MLLT3, MNX1, MSX1, MSX2,* and *NRIP1*). Of the transition genes several have been reported to be involved in axial elongation (*GREB1* (*60*), MNX1 (*61*), *MSX1* and *MSX2* (*62*), while others are associated with neural development (FABP5 (*63*) and EMX2 (*64*)). We also included the three genes *F2RL1*, *CMTM8*, and *MLLT3* which show distinct expression patterns within the CLE.

**Table 2:**
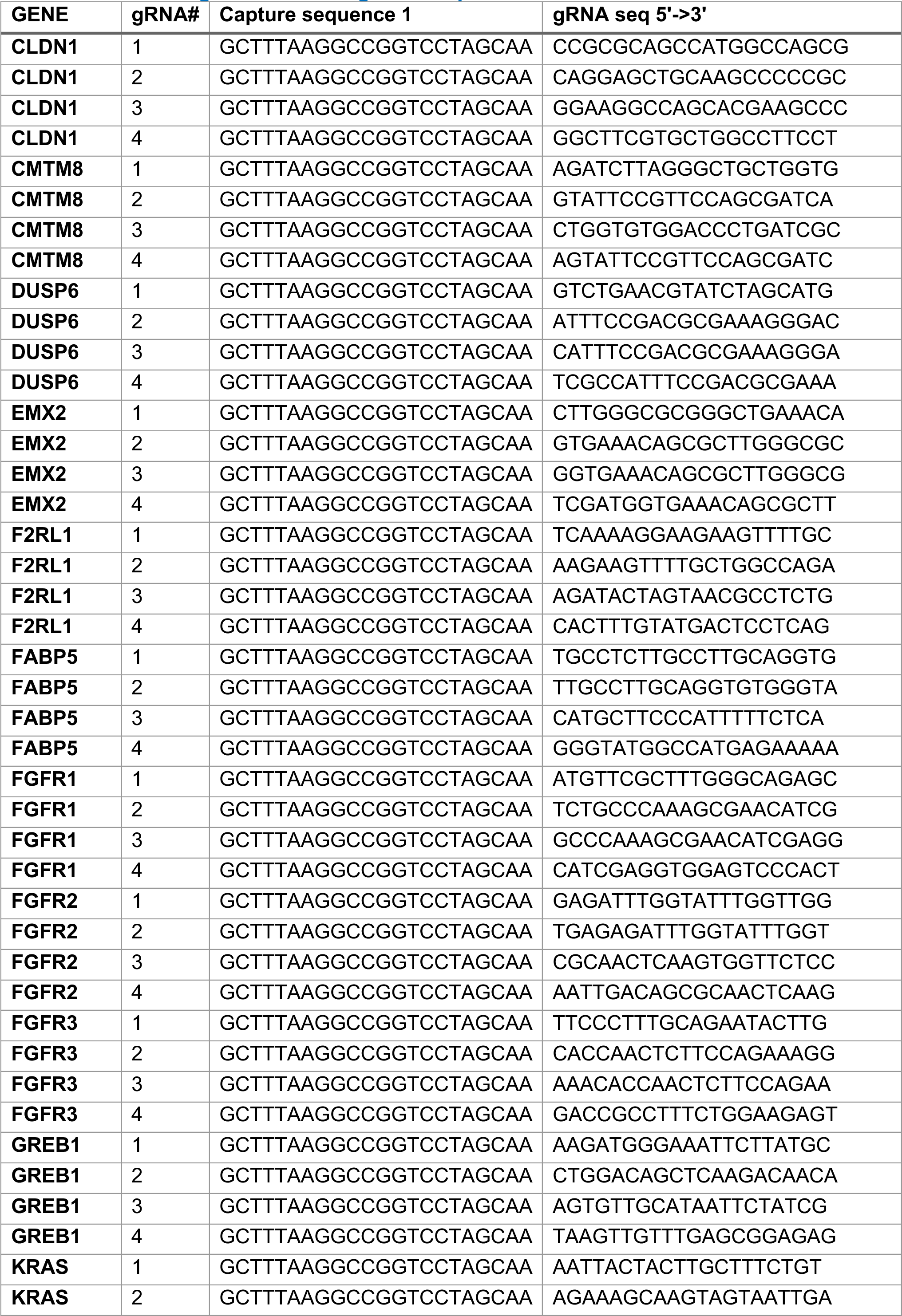
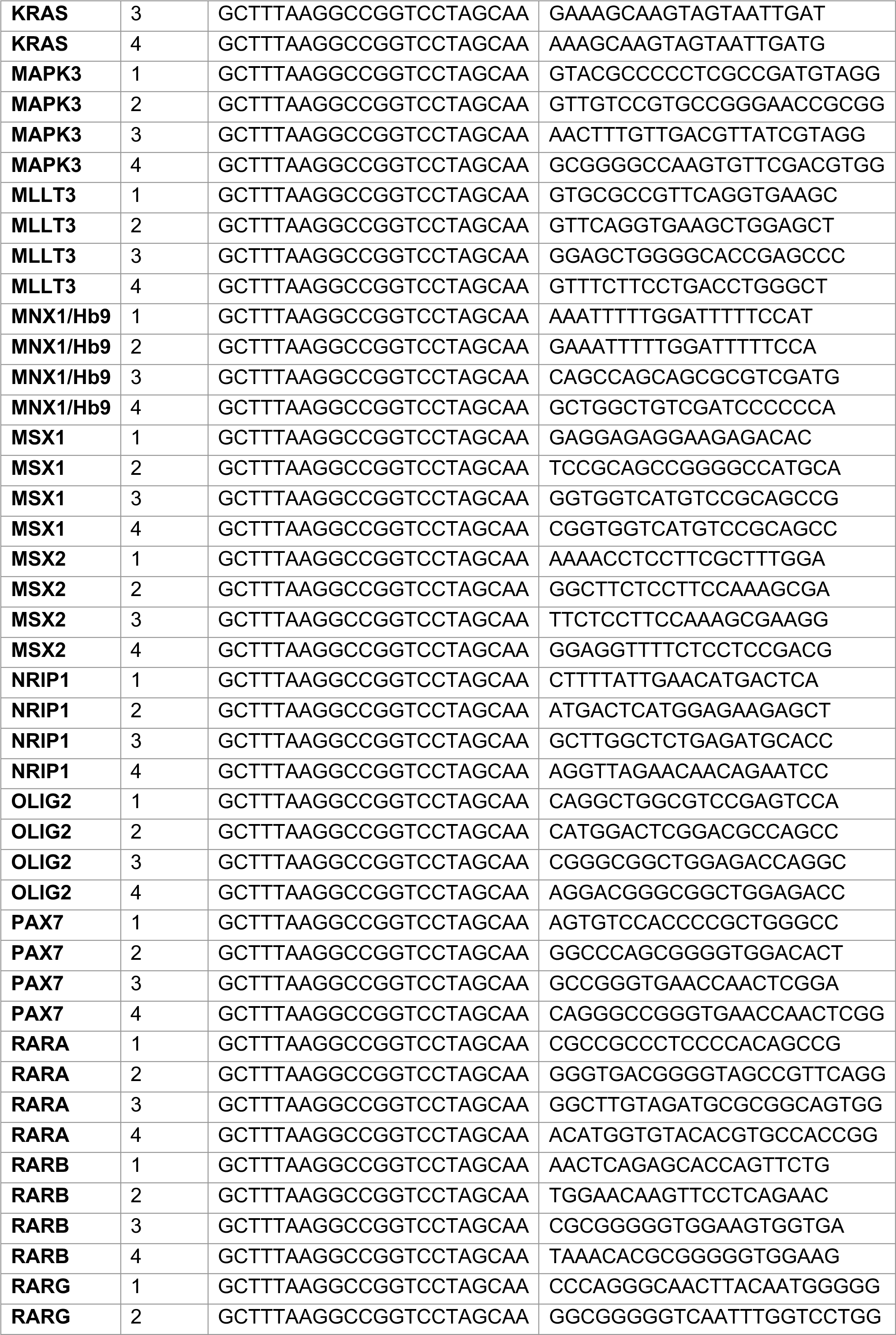
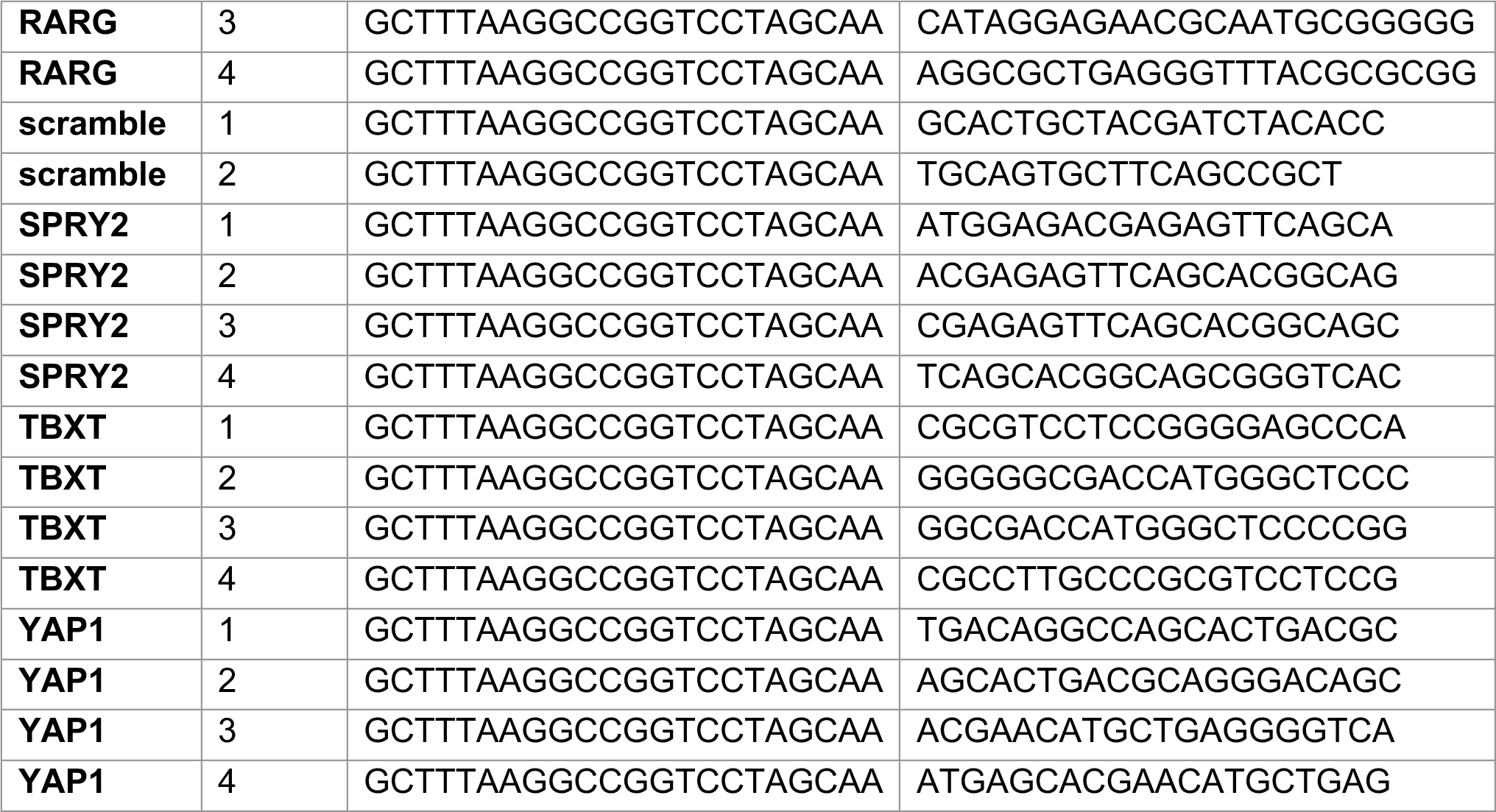
List of Target Genes and gRNA sequences.

We then performed targeted electroporation of the plasmid pool to the CLE of HH9 chick embryos (Figure 3A, left: n = 4 replicates of 10 embryos/replicate) and tracked the outcome of individual gRNAs by examining the fates of cells 24h post-electroporation by enriching Cas9+ cells via FACS and single cell RNA sequencing (Figure 3A - right). We hypothesized that a knockout could result in cell death (loss of guide detection), increase cell proliferation (increased guide detection), changes in cell identity (increase or loss of guide detection in particular fates), or misregulate gene expression within a cell.

**Figure 3:**
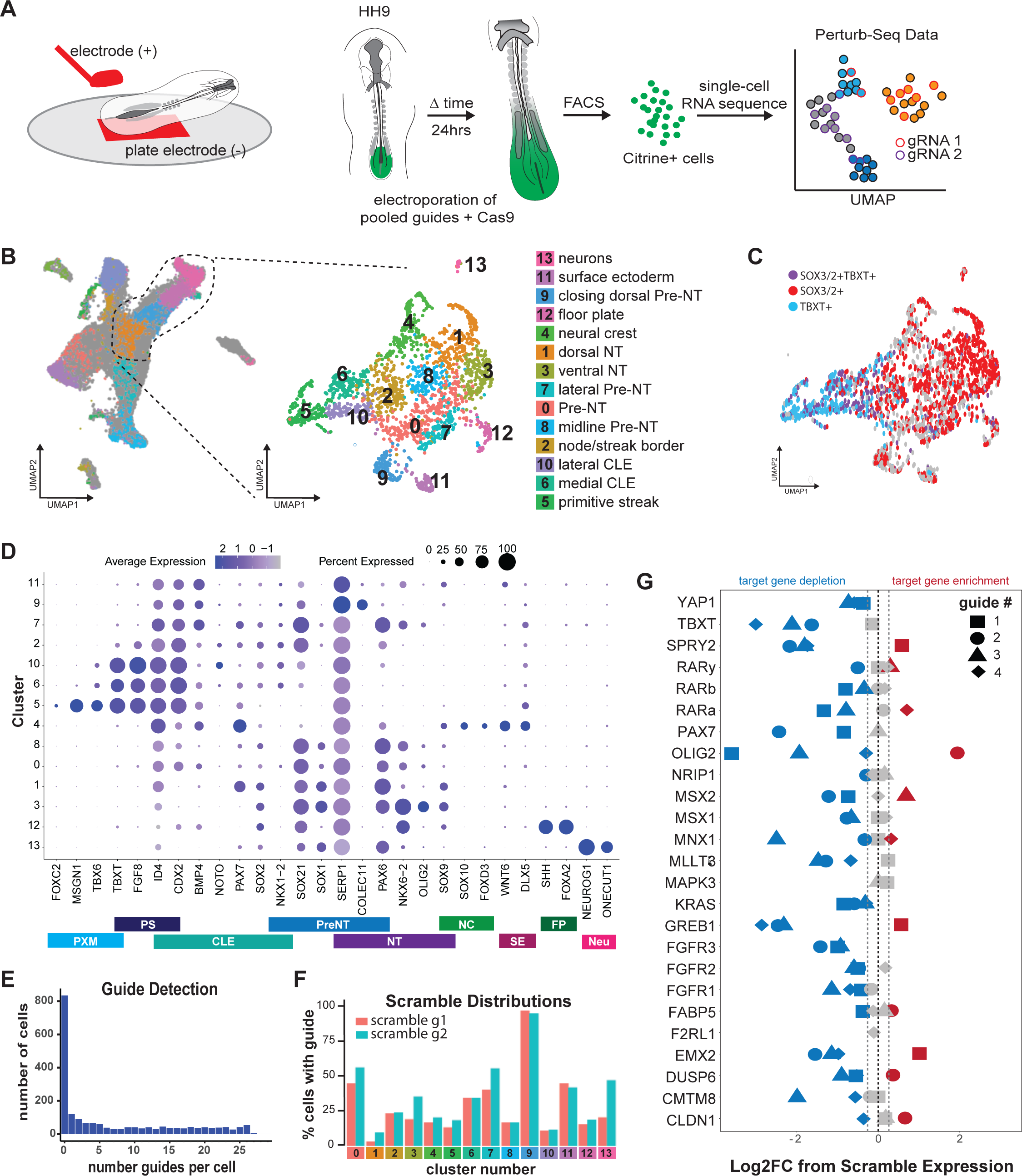
Pooled in vivo screen in chick embryos. A) Schematic of electroporation and screening workflow. HH8 chicken embryos were electroporated with CRISPR plasmids ex ovo, allowed to develop for 24h and subsequently enriched for electroporation via FACS of Citrine positive cells. Cells were profiled by single-cell RNA sequencing. B) UMAP projection of single cell RNA sequencing data into wildtype reference map (left) and subsequent selection and re-clustering of primitive streak, CLE and neural tube (right). C) UMAP showing NMPs in screen dataset marked by co-expression of *SOX2/3* and *TBXT*. D) Dot plot representing gene expression across screen dataset of lineage markers. PXM – paraxial mesoderm, PS – primitive streak, CLE – caudal lateral epiblast, PreNT – preneural tube, NT – neural tube, NC – neural crest, FP – floor plate, Neu – maturing neurons. E) Number of guide barcode UMI counts detected per cell across screen dataset after filtering for ambient RNA amplification. F) Percent of cells that display a scramble gRNA barcode distributed across clusters of screen dataset. G) Differential expression of genes targeted by each guide in cells that received said gRNA (blue: reduction in expression compared to scramble, red: increase compared to scramble, grey: no change in expression, dashed line marks 5% false discover rate (FDR)).

We projected the single-cell transcriptomes of targeted cells onto the wildtype dataset, subsetted cells classified as primitive streak, CLE, and neural tube, and then reclustered (Figure 3B). The resulting 13 clusters were annotated using the same markers as the wildtype dataset (Figure3D, S4A). Similar to the wildtype dataset, we identified, *SOX2, SOX3* and *TBXT* expressing NMPs, marking the CLE (Figure 3C). We additionally observed a cluster of *NEUROG1*+ neurons, a cluster that expressed *WNT6*, *DLX5* and *SERP1* positive surface ectoderm, and *BMP4+COLEC11+* cells which is consistent with the closing the dorsal neural tube (Figure 3C,D, Figure S4A).

To determine knockout phenotypes, we used gRNA capture sequences to identify perturbations present in individual cells. Following stringent filtering steps excluding amplification below two standard deviations of the mean as ambient RNA amplification (Figure S5B), we found 0-29 detectable guides per cell (Figure 3E). Importantly, both scramble gRNAs were detected in the dataset and of the 102 guides used only 5 were undetected in the final dataset: PAX7 guide 4, MAPK3 guide 4, and F2RL1 guides 1-3.

It has been previously reported that individual cells receiving multiple guides may have confounding effects of multiple knockouts. However, trends for a particular perturbation would be maintained across all cells that had received that guide (*34*). To test this, the distribution of cells with either scramble guide regardless of multiple knockouts was compared under the assumption that the distribution of the two scramble guides would resemble one another. Despite some heterogeneity in guide distribution across clusters, the two scramble guides were equally represented (by ANOVA and subsequent Tukey tests), consistent with previously published results of pooled screens *in vitro* where multiple guides per cell were detected. We therefore used the average of the two scrambles to represent expected representation of gRNA in each of the clusters.

As one of the *OLIG2* guides did not effectively deplete Olig2 (Figure 2C), assumed guides targeting other genes would have a similar trend. Thus, we defined “successful” guides as those reducing target gene expression by comparing cells that received a target gRNA to those that received a scramble gRNA (Figure 3G). This stringent criterion yielded 2-3 “successful” guides/gene on average. Notably, the sole detected *F2RL1* guide (guide 4) was “unsuccessful,” which in addition to the dropout of F2RL1 gRNAs 1-3 suggests that this gene may be necessary for cell survival in the CLE. While possible dropout could be a result of lower concentration of *F2RL1* guide, the hypothesized cell death with *F2RL1* knockout supports previous evidence that loss of *F2RL1* is partially embryonic lethal (*65*) and loss of both *F2RL1* (Proteinase-Activated Receptor 2: *PAR-2*) and *F2R* (*PAR-1)* result in neural tube defects (*66*). Overall, we were able to develop and confirm the successful targeting of multiple genes in a pooled manner within single embryos.

### Guide enrichment highlights importance of FGF and RA signalling in epiblast to neural fate transition

As cells in the CLE generate multiple cell types, we examined whether specific gRNAs altered the distribution of cell types by assessing guide enrichment compared to scramble controls. First, we used chi-squared tests to compare guide representation in each cluster to both the scramble guide 1 and the average of all guides (Figure 4A). Only *YAP1* gRNA containing cells did not differ from scramble controls, indicating that *YAP1* perturbation had no observed effect on lineage fate proportions. However, Chi-squared tests are sensitive to populations with low numbers, we sought to confirm the Chi-squared results using a Kullback-Leibler Divergence (KLD) test, comparing the distribution of targeting guides across clusters to the average scramble distribution (Figure S6A). Of the 25 gene knockouts analysed (excluding *F2RL1* and *MAPK3* with no successful guides from previous filtering steps), only *YAP1* knockout did not significantly differ from scramble, corroborating the Chi-squared analysis.

**Figure 4:**
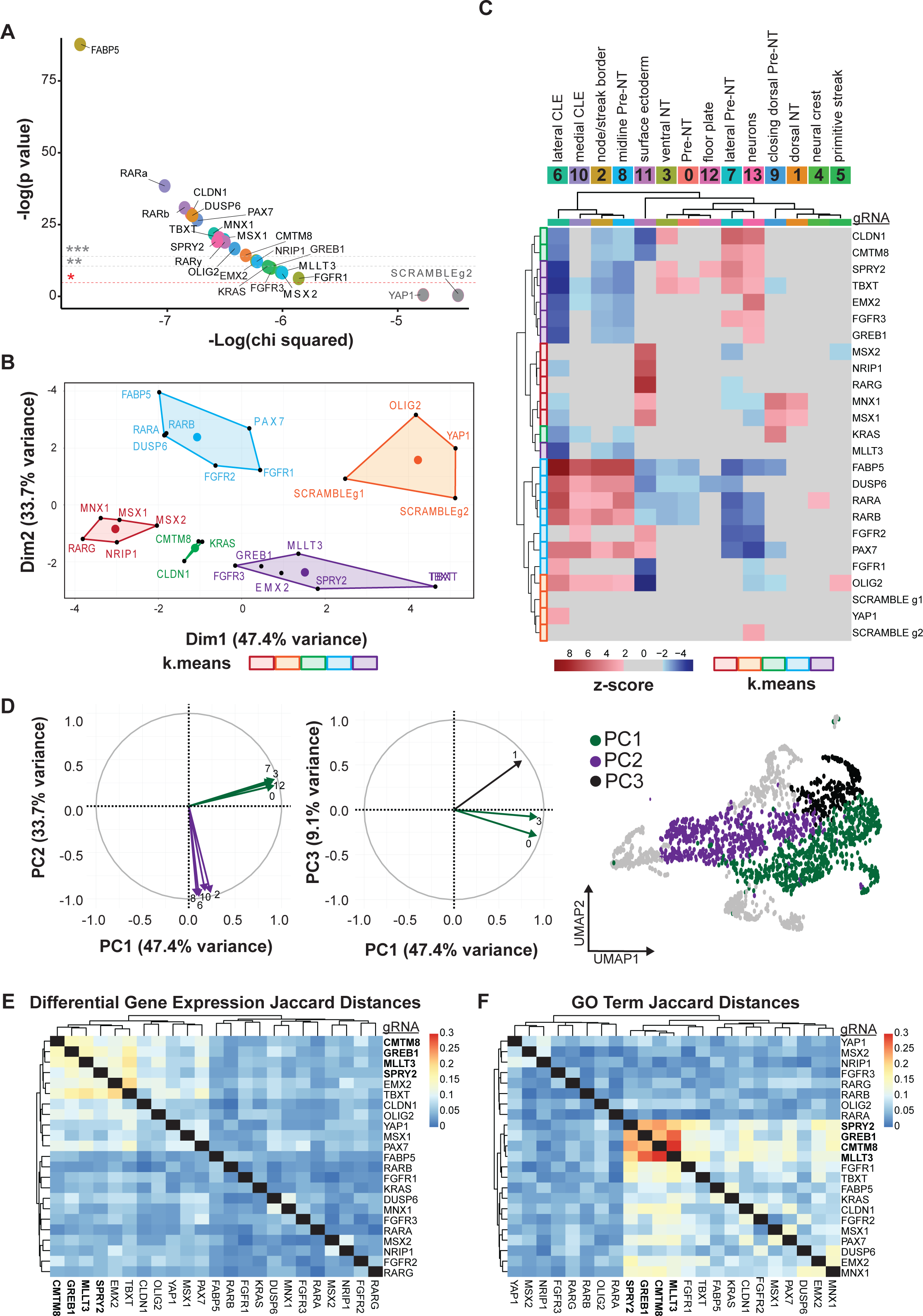
Genes affecting transition from epiblast to neural tube.

A) Chi-squared tests comparing distribution of gRNA containing cells to the average across all gRNAs and to the distribution of scramble 1. Non-significant gRNAs are labelled in grey. (* = p <0.05, ** = p<0.001, *** = p< 0.0001). B) K-means clustering of gRNA enrichment scores for each gRNA. Cluster number was determined by elbow plot of the intra-cluster distances. C) Left: Heatmap of gRNA containing cell enrichment compared to scramble within each cluster of the dataset, marked by chi-squared test z-scores above 2 and below −2. D) Radial plot of principal component analysis of enrichment between knockouts depicting which clusters are responsible for each component. Right: schematic of screen dataset UMAP colored by drivers of each principal component (PC1 – purple, PC2 – green, PC3 – grey). E) Heatmap of calculated Jaccard distances between each set of identified differentially regulated genes post knockout. F) Heatmap of Jaccard distances based on the list of Gene Ontology Terms generated from the differentially regulated genes of each knockout.

We then calculated gRNA enrichment or depletion relative to scramble for each cluster (Figure 4C). K-means clustering segregated gRNA containing cells into 5 classes based on enrichment/depletion patterns (Figure 4B,C). The two scramble gRNAs and *YAP1* gRNAs cluster together. *OLIG2* gRNA enrichment pattern also clustered with the scramble and *YAP1* gRNAs, this might reflect the earlier stage of development and the limited number of neural progenitors in the screen dataset compared to the pilot experiments. Principal Component Analysis (PCA) revealed the largest variation in enrichment/depletion (47.4%) was driven by depletion in clusters representing ventral neural tube progenitors and pre-neural tube (Dim1, Figure 4D). PC2 (33.7% variation) represented enrichment/depletion in clusters corresponding to the CLE, node-streak border, and medial pre-neural tube (Dim2, Figure 4D). PC3 (9.1% variance) involved clusters representing the dorsal neural tube (Figure Dim3, Figure 4D). This cluster analysis reveals that many perturbations either affected cells’ ability to become neural (PCA1) or their ability to remain in the CLE and maintain stemness.

Following the overarching trend of tight regulation of CLE exit, gRNAs targeting FGF receptors 1 and 2, the ERK/FGF inhibitor *DUSP6*, and retinoic acid receptors RARα and RARβ clustered together and showed enrichment of cells in clusters containing axial progenitors and depletion in those containing neural fates. FGF and retinoic acid pathways regulate the establishment of the neuroepithelium and the exit of cells from the CLE with opposing roles in maintaining stemness (FGF) and promoting differentiation (RA)(*4, 27*). The co-enrichment of FGF receptor and RAR knockouts in the CLE is consistent with the necessity of these pathways for the CLE-to-neural transition. Additionally, *FGFR3* and the FGF inhibitor *SPRY2* were depleted from the CLE, clustering with steroid signalling or WNT signalling (*GREB1, TBXT, MLLT3 and EMX2*). This suggests roles for these pathways in maintaining CLE populations. Overall, examining changes in knockout cell ratios identified two main contributors to acquisition of cell fate: the ability to differentiate to neural fates and an increase/depletion from the progenitor pool of the caudal epiblast.

### *CMTM8*, *GREB1*, *MLLT3*, and *SPRY2* perturbation causes similar down-stream gene misregulation

As the perturbation of transition-associated genes affected the CLE-to-neural fate commitment, we then investigated disrupted gene expression that might lead to this change in differentiation. To test this, we first examined the full set of gRNAs, regardless of enrichment/depletion distributions compared to scramble controls. To focus on neural fate decisions, we analyzed changes in the expression of the top 500 cESFW-selected genes per cluster from our wildtype dataset (2505 total genes). Using the package scMAGeCK, designed to identify gene regulation in pooled CRISPR screens (*67*), we generated lists of differentially expressed genes resulting from each gRNA and performed K-means clustering to group knockouts with similar effects. There was a high level of variability between the differentially expressed genes for each knockout, exemplified by the near-linear intra-cluster distance loss (Figure S6C). However, calculating the Jaccard distances between the misregulated genes for each gRNA (Figure 4E) revealed a group of consistently misregulated genes associated with *CMTM8, GREB1, MLLT3, SPRY2, EMX2,* and *TBXT* gRNAs (maximum Jaccard distance of 0.2, indicating 20% overlap).

As the screen also highlighted the regulation of WNT, FGF, and RA signalling pathways, we asked whether the same biological pathways and processes were affected with different gRNAs. Examining Gene Ontology (GO) and KEGG pathway categories for the biological processes that were affected by the differentially expressed genes of each gRNA, revealed 30-40% overlap (Figure 4F, Figure S6B). The GO term clustering highlighted the importance of the FGF and RA signalling pathways in establishing trunk lineages. Jaccard distances of knockouts associated with RA and FGF/ERK hierarchically clustered separately (Figure 4F - left and right respectively). These two signalling pathways have been repeatedly implicated in generating the diversity of cell types within the spinal cord and neighbouring somites (*27, 68, 69*) and our screen results corroborate these findings. Moreover, perturbations to the previously highlighted group of gRNAs (*SPRY2*, *GREB1*, *CMTM8*, and *MLLT3*) displayed the highest GO term and KEGG pathway overlap (20-30%). This is consistent with *SPRY2’s* role in fine-tuning FGF signalling in the trunk (*70*) and loss of *GREB1* affecting downstream WNT signalling and trunk elongation (*60*). Hoever, the roles of *CMTM8* and *MLLT3* influence the regulation of trunk development has yet to be explored.

### *MLLT3* gRNAs Affect Multiple Trunk Signalling Axes

From our gRNA enrichment analysis, *SPRY2, GREB1, CMTM8, MLLT3* gRNAs were depleted in the CLE. To investigate this further, we examined overlapping misregulated genes between the perturbations (Figure S7A). All four gRNAs affected Wnt pathway genes (*WNT5A, TBXT*), decreased *FGF8*, and increased *FGFR3* expression, highlighting a potential role in self-renewal of progenitors within cells of the CLE. Examining the parent GO terms between gRNAs (Figure 5A) indicated processes such as cell division, migration, neuronal differentiation, pattern specification, retinoic acid signalling, regulation of neuronal death, and Wnt signalling (Figure 5B). *MLLT3* perturbation alone was associated with both Wnt signalling and RA signalling, connecting *MLLT3* to two pathways regulating stemness in the CLE (Wnt) and neural differentiation (RA). We therefore focused on *MLLT3* to elucidate its role within the CLE and neural tube.

**Figure 5:**
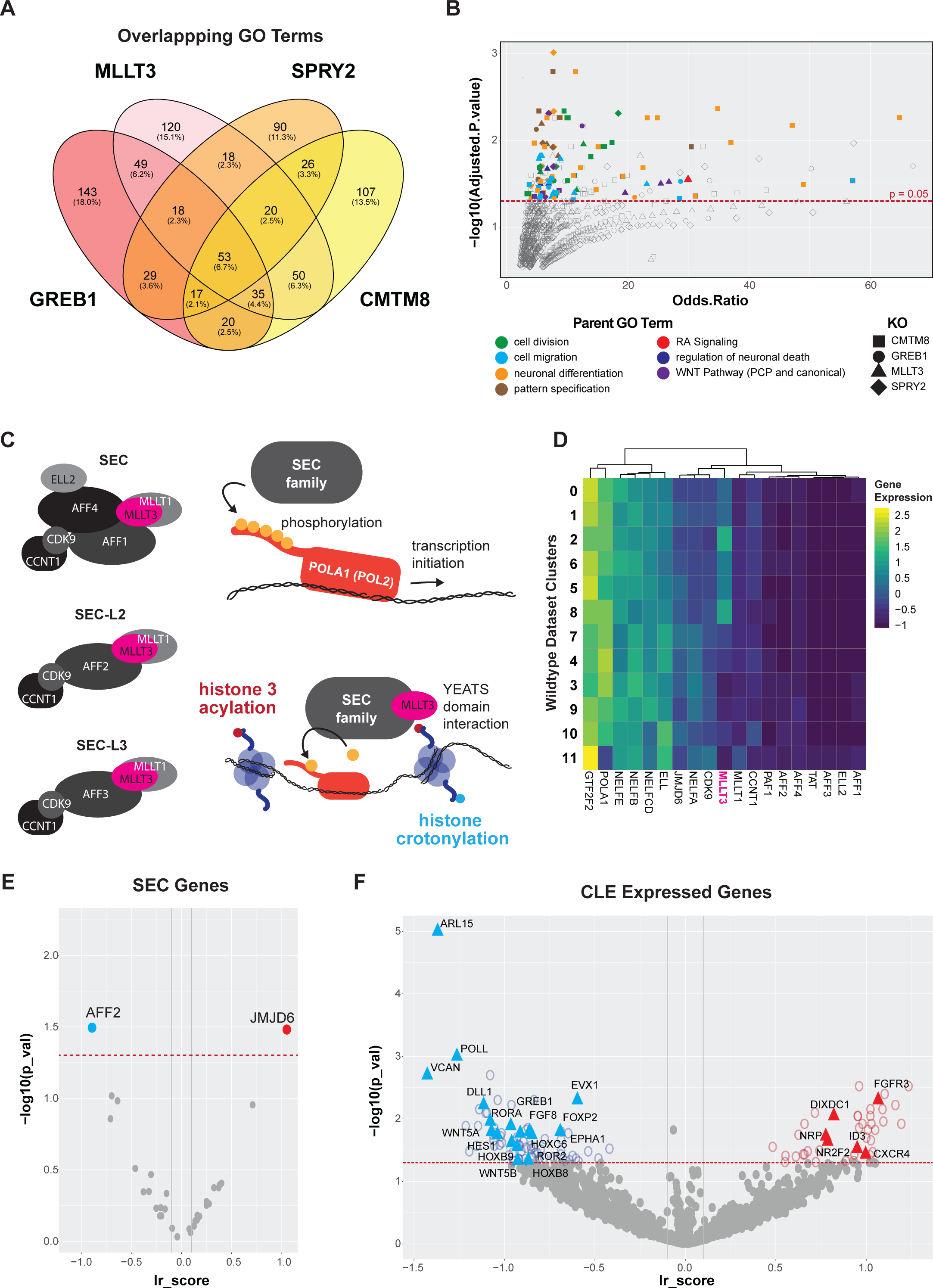
Targeting MLLT3 disrupts Wnt PCP and RA signalling in the CLE.

A) Venn diagram showing the overlapping percentages of Gene Ontology (GO) terms between *MLLT3, SPRY2, GREB1*, and *CMTM8*. B) Significance of GO terms where color marks the parent classification of GO term: green – cell division, light blue – cell migration, yellow – neuronal differentiation, brown – pattern specification, red – Retinoic Acid signalling, navy blue – regulation of neuronal cell death, purple – Wnt signalling, and shape marks the knockout. (red line indicates adjusted p = 0.05 by Fisher’s exact test). C) Schematic of mammalian super elongation complex family members (SEC, SEC-L2, SEC-L3) and the phosphorylation of Poll (*POLA1* in chicken). D) Heat map indicating the detected members of the SEC family and related complexes in the wildtype reference single-cell RNA sequencing dataset. *MLLT3* highlighted in pink. E) Differential regulation of SEC complex family members. (lr score is a corrected read out for differential gene expression (*67*), red line indicated p = 0.05 FDR). F) Differentially regulated genes post *MLLT3* perturbation, from the Entropy Sort Score ranked gene list, (lr score is a corrected read out for differential gene expression (*67*), red line indicated p = 0.05 FDR).

*MLLT3* is a YEATS family member and super elongation complex (SEC) component (*59–62*). It participates in the three mammalian forms of SEC (Figure 5C) to facilitate rapid gene induction by interacting with acetylated/crotonylated histone lysines (*71–74*). *MLLT3* has primarily been studied in cancer, where its truncation and fusion with *MLL1* leads to aberrant H3K79 methylation, dysregulating the JNK (*75*) and Wnt pathways (*76*). In developmental contexts, *MLLT3* has been implicated in vertebrae malformations (*77*) hematopoietic stem cell self-renewal (*78*), and hematopoietic lineage decisions (*79*). MLLT3 has also been shown to bind retinoic acid receptor α (RARα) and activate neurogenic programs in vitro in mammalian cells (*80*), revealing a possible role in regulating a range of signalling pathways.

Because little is known about the SEC complex members in chicken, we first examined the reference wildtype single cell RNA sequencing dataset to determine whether the components of mammalian SEC complexes were expressed (Figure 5D). We found SEC-L2 and SEC-L3 were expressed in the epiblast and neural tube indicating possible transcriptional regulation by all three complexes in chicken similar to mammals. Of the expressed components, only *MLLT3* was differentially expressed across the CLE to neural lineage trajectory and as such the only SEC component that was selected by the previous Entropy Sorting analysis as a hit for lineage regulation. Further, of the proteins associated with the SEC, the demethylase *JMJD6* (*81*) and the scaffold protein *AFF2* were up and downregulated, respectively with *MLLT3* perturbation (Figure 5E), indicating a loss or change in composition of the SEC complex formation in the absence of *MLLT3*.

To investigate the effect of *MLLT3* perturbation on gene expression, we examined differentially expressed genes in the *MLLT3* gRNA containing cells compared to scramble (Figure 5F). Genes associated with RA signalling and neural differentiation such as *RORA, EVX1*, and *FOXP2* (*26, 82, 83*) were reduced in *MLLT3*-perturbed cells, while genes repressed by RA signalling, such as *NR2F2* (*84, 85*), were upregulated. Additionally, Wnt PCP pathway genes important for caudal epiblast fate (*WNT5A, WNT5B, ROR2*) (*86*) and Notch signalling genes involved in neural fate acquisition (*DLL1, HES1*) were downregulated. The misregulation of multiple pathways with opposing functions, positions MLLT3 as a possible connection between CLE stem cell maintenance and acquisition of neural fate. This prompted us to investigate further the role of *MLLT3* in epiblast and neural tube lineage transitions.

### MLLT3 regulates neural tube fate acquisition through retinoic acid receptor binding

To investigate *MLLT3*’s role without the potential confounding factors of a pooled screen, we repeated individual *MLLT3* targeting in the CLE of HH9 chick embryos. Consistent with the screen results, after 24h MLLT3 gRNAs caused both reduction in MLLT3 expression (Figure 6A, S8A) as well as a disruption of the *MLLT3* expression domain in the tail bud (Figure S8B). Further, MLLT3 KO embryos displayed kinked spinal cords with higher tortuosity (Figure 6B). Additionally, we observed a reduction of *MLLT3* gRNA containing cells in the neural tube and tail bud (Figure 6A:a’-d’). To test whether this reduction was cell autonomous, we co-electroporated the MLLT3 CRISPR targeting constructs with a non-targeting Ef1a:NLS-mScarlet construct (Figure 6B). Here the mScarlet cells serve as a within embryo control that should maintain their ability to differentiate to neural tube lineages. Whole-mount staining and flow cytometry revealed depletion of Citrine+ cells from the neural tube in the *MLLT3* gRNA transfected condition, while control mScarlet+ cells in the same embryo were present in the neural tube and maintained SOX2 expression (Figure 6C,D, Figure S8C). Importantly, the total Citrine:mScarlet ratio of cells remained consistent between scramble and *MLLT3* gRNA conditions (Figure S8D), indicating *MLLT3* disruption does not cause cell death, but rather affects cell fate and results in depletion from neural tube lineages.

**Figure 6:**
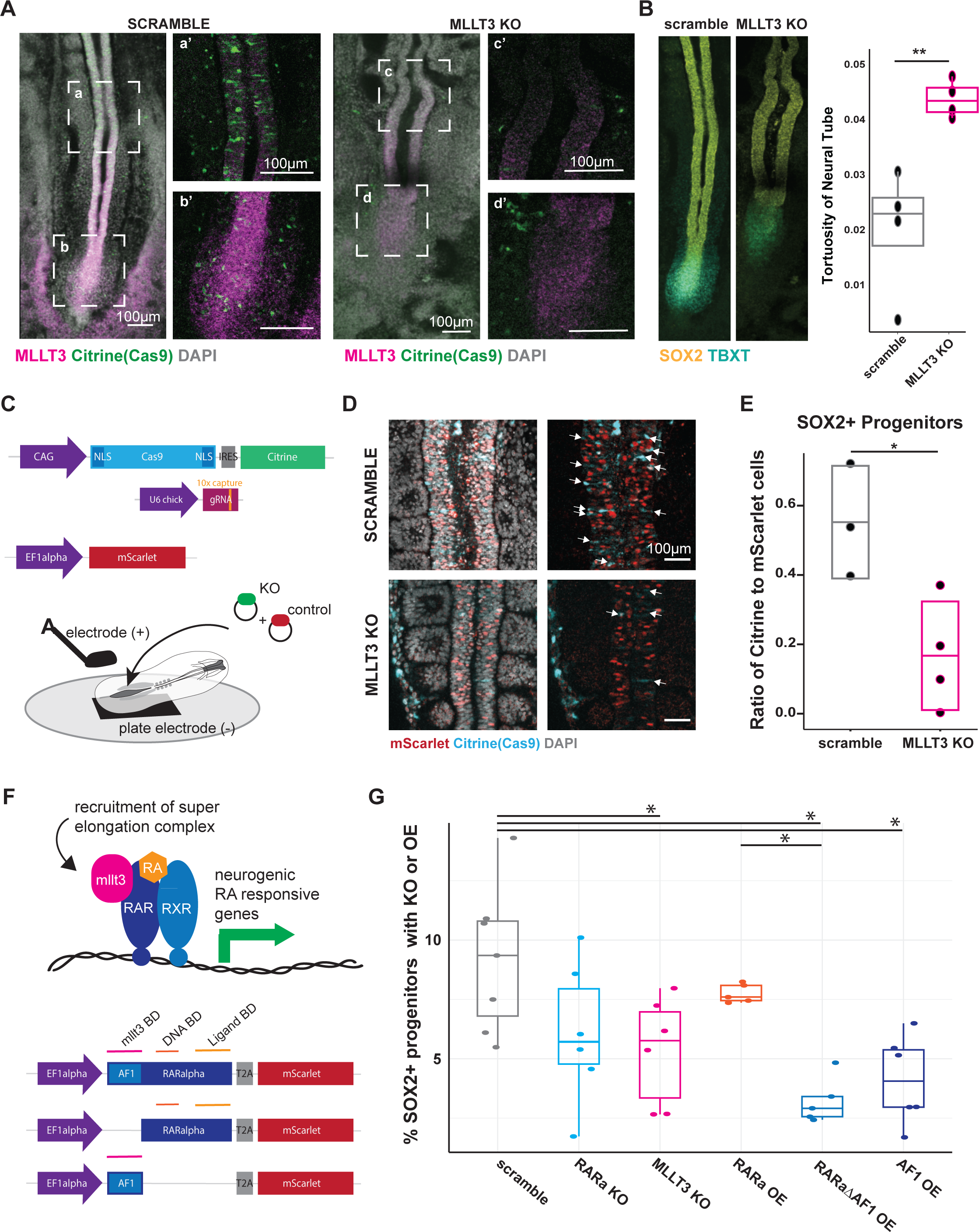
MLLT3 regulates neural progenitor formation through RARα interaction.

A) Fluorescence images of hybridization chain reaction for *SOX2, TBXT, MLLT3*, and reporter fluorescence of Citrine (marking Cas9 containing cells) for scramble (left) and MLLT3 (right) gRNA transfected embryos. B) *LEFT:* paired HCR images from 6A depicting *SOX2* and *TBXT* epression in embryos post perturbation. *RIGHT:* Quantification of changes in tortuosity (perimeter/area) of neural tube based on masks of *SOX2* expression in embryos post perturbation. (n = 4 embryos per condition, ** indicates p<0.01 via t-test). C) Schematic of two-colour electroporation of caudal epiblast of HH9 embryos. D) Immunofluorescence images showing loss of Citrine (Cas9) in *MLLT3* gRNA neural tubes and maintenance of mScarlet electroporation control. E) Flow cytometry of SOX2+ neural progenitors in scramble versus *MLLT3* KO embryos where the y-axis depicts the ratio of mScarlet to Citrine positive cells (* indicates p<0.05 by t-test, n = 4-5 embryos per condition). F) *TOP:* Schematic of possible action of Mllt3 to regulate RA signalling response. *BOTTOM:* Schematic of constructs of RARα indicating the Mllt3 binding domain (Mllt3 BD), the DNA binding domain (DNA BD) and the ligand binding domain (ligand BD), visualized by mScarlet. G) Quantification of flow cytometry analysis of SOX2+ cells in embryos that have been electroporated with the indicated CRISPR construct or over expression constructs (* indicated p<0.05 by ANOVA and subsequent Tukey tests, n = 5-7 embryos per condition).

As *MLLT3* interacts with RARα and contributes to neural differentiation in vitro (*80*)(Figure 6E), we investigated whether the depletion of *MLLT3* KO cells in the neural tube is due to RARα binding and presumed elongation complex recruitment to neural genes. We compared caudal epiblast-specific *MLLT3* perturbation to the forced expression of mutant RARα proteins: one lacking the *MLLT3* binding domain (ΔAF1), and one of the AF1 binding domain alone (Figure 6E). Flow cytometry analysis of SOX2 expressing neural tube progenitors reconfirmed a decrease in neural progenitors compared to scramble when targeting *MLLT3*. Depletion via CRISPR gRNA of RARα or wildtype RARα overexpression had no significant effect, potentially due to compensatory mechanisms. However, forced expression of both RARα mutant constructs phenocopied *MLLT3* knockout with reduced SOX2+ neural progenitors. Together, this provides evidence of a previously undocumented role of *MLLT3* in the epiblast-to-neural transition via interaction with the retinoic acid pathway and downstream genes.

## DISCUSSION

The caudal epiblast of vertebrate embryos contains populations of cells that generate the tissues of the trunk during axis elongation. The coordinated generation of these tissues depends on the signalling dynamics to determine the rate at which cells differentiate and the proportions that are allocated to specific lineages. How this is organized, and the gene regulatory mechanisms responsible for the process remain poorly defined. Mapping gene expression in the CLE called attention to several factors that might have functional roles in the elaboration of trunk tissues. To test this, we establish methods for multiplexed in vivo CRISPR screening in chick embryos by incorporating capture sequences to gRNAs in a chick-specific CRISPR system (*48, 49*). The screen successfully enriched for regulators of cell fate decisions, evidenced by the clustering of cells containing gRNAs targeting specific signalling pathways and gene expression programmes associated with neural fate specification. Further, the screen identified a previously undocumented role for MLLT3, a component of the super elongation complex, in the acquisition of neural cell fate. This highlights the power of combining transcriptomics with functional genomics. More generally, the validation of the CRISPR system for functional in vivo assays, paired with the historically well-characterized embryology of the chick and the potential for regionally targeted electroporation (*87, 88*), opens up many possibilities for interrogating gene regulatory mechanisms in vivo in a highly tractable system.

The screen highlighted the importance of RA, WNT and FGF signalling pathways in regulating the epiblast-to-neural transition. Perturbation of FGF receptors, the ERK inhibitor DUSP6, and Retinoic Acid Receptors resulted in the accumulation of cells in the caudal epiblast and depletion of ventral neural fates (Figure 4C), consistent with the roles of FGF and RA in self-renewal and differentiation respectively. FGF signaling is needed to trigger early neural differentiation (*25*), emphasizing the need of precise regulation of signalling. Conversely, targeting the FGF inhibitor SPRY2 and WNT pathway components depleted the epiblast progenitor pool, suggesting these genes are critical for CLE maintenance. These results confirm and extend our understanding of the dynamic interplay between these signalling cascades in controlling fate decisions in the CLE.

Further, the screen identified a previously unobserved role for the gene MLLT3 as a key regulator for the initiation of the neural gene expression programme. An epigenetic reader of histone lysine acetylation and crotonylation, MLLT3 contributes to gene expression as part of the super elongation complex (*71, 74, 89*). In development, MLLT3 has been studied in mammalian hematopoiesis (*90*), and further maintains stemness in adult hematopoiesis and its overexpression expands and increases engraftment of transplanted hematopoietic stem cells (*78*). Our analysis indicated that loss of MLLT3 disrupted caudal epiblast fate maintenance (Figure 3C), possibly prematurely depleting this stem cell reservoir and compromising neural tube formation (Figure 6). Transcriptomic analysis of MLLT3 knockout cells suggested dysregulation of multiple signalling pathway components implicated in neural induction, including retinoic acid, WNT, and Notch signalling cascades. The specific factors misregulated, such as decreased expression of RA-induced neurogenic genes (RORA, ROR2, EVX1, FOXP2) and components of the WNT/PCP and Notch pathways (WNT5A/B, DLL1, HES1), provide a possible mechanistic explanation for the impaired neural fate acquisition upon MLLT3 perturbation.

MLLT3 has been shown to interact with a variety of transcription factors to influence differentiation programmes. For example, binding to GATA1 directs erythroid and megakaryocyte differentiation (*79*), while interacting with TET2 drives cortical neuron generation (*91*). Direct binding of MLLT3 to Retinoic Acid Receptors recruits the super elongation complex to rapidly induce neural programmes (*80*). Moreover, a mutant form of RXRα disrupts proliferation in murine KMT2A-MLLT3 leukemia cells (*92*). Consistent with a relationship between MLLT3 and RA signalling, our results from complementary approaches, indicate that disrupting the ability of RARα to interact with MLLT3 phenocopied the neural tube defects observed with perturbing *MLLT3*. These results implicate MLLT3 as a critical cofactor that facilitates the gene expression changes driven by retinoic acid signalling during neural induction. MLLT3 serves as an example of how the precise spatial and temporal control of transcription can drive rapid shifts in gene expression in response to signalling cues, a regulatory strategy likely employed broadly during embryogenesis. Examining the interplay between signal-responsive transcription factors and elongation complexes will be critical for understanding fate specification across contexts.

While this focused approach successfully identified MLLT3, expanding the size of our gRNA library could uncover additional regulators and provide a more comprehensive view of the epiblast-to-neural transition. The pooled screening approach results in individual cells receiving multiple guides targeting different genes. Although this did not appear to affect the average distributions of knockout cells in our study, and has been shown previously to be an effective screening method (*34*), it may mask subtle effects on gene networks or lineage commitment. Furthermore, considering only those guides that decreased the level of the targeted transcript may unnecessarily exclude some data from the analysis. Although we demonstrate loss of neural progenitors with mutant RARα constructs directed at the AF1 binding domain, this domain interacts with co-factors other than MLLT3 and disruption of these interactions might contribute to the observed effects. For example, BRD4 interacts with the AF1 domain (*80*). Though expression of BRD4 was not detected in either the wildtype or the screen datasets, it indicates the possible action of other co-regulators in this process.

In summary, the application of an in vivo pooled CRISPR screening strategy in chick embryos has uncovered an unexpected role for the super elongation factor component MLLT3 in the differentiation of neural cell fates from caudal epiblast cells. The findings provide new mechanistic insight into the gene regulatory logic underlying neural tube formation and demonstrates the power of this screening platform to interrogate developmental processes at single-cell resolution in an embryonic model system. With further optimization, this approach holds promise for the detailed dissection of gene regulatory mechanisms across diverse tissue contexts.

## ACKNOWLEDGEMENTS

We thank Giulia Boezio and Despina Stamataki for help in establishing chicken protocols, Tatiane Nakamura Kanno for the ex ovo culture protocol, Tatjana Sauka-Spengler and Ruth Williams for the CRISPR plasmids, and Philippe Lefebvre for the plasmids containing the RARα mutant sequences. We are grateful to members of the Briscoe Lab for their constructive feedback on the manuscript as well as Gunes Taylor, Naomi Moris, and Tatiane Nakamura Kanno. We thank the following Science Technology Platforms at the Francis Crick Institute for their expertise and assistance: Advanced Sequencing STP, Making Lab STP, Advanced Light Microscopy STP, Biological Research Facility, Experimental Histopathology, and Flow Cytometry STP. This work was supported by the Francis Crick Institute which receives its core funding from Cancer Research UK (CC001051), the UK Medical Research Council (CC001051), and Wellcome (CC001051); by the European Research Council under European Union (EU) Horizon 2020 research and innovation program grant 742138; and by Wellcome (226633/Z/22/Z; 220379/Z/20/Z). ARGL was supported by EMBO ALTF (149-2020).

## AUTHOR CONTRIBUTIONS

ARGL and JB conceived and designed the project, interpreted the data, and wrote the manuscript. ARGL characterized embryos via HCR and immunofluorescence staining, cloned the CRISPR and over expression plasmids, performed single-cell RNA sequencing and all chick electroporation experiments. TR performed quantification of HCR images. AR applied Entropy Sorting analysis to the wildtype dataset. TR and AR provided editorial contribution to the manuscript.

## METHODS

### Egg incubation, embryo staging, electroporation and ex ovo culture

Eggs obtained from Henry Stewart & Co. Ltd were incubated for 36 hours at 38°C to generate Hamburger Hamilton stage (HH) 9 embryos with 7 somites (*93*). Two types of electroporation were conducted: in ovo and ex ovo using an Electro Square Porator ECM830. All plasmid mixes were injected via a pulled glass capillary (Harvard Apparatus, EC1 64-0766) and contained Sybr green (ThermoFisher, S7563), PBS, the Cas9 plasmid at a concentration of [500ng/ul] and the pool of guide plasmids at a concentration of [50ng/ul] per individual guide plasmid.

In ovo electroporation involved injection of plasmid mix through the head of the HH9 embryo and down the neural tube where electrodes were placed on either side of the neural tube (pulse regime - 30V, 50ms length, 3 pulses, 200ms interval). The egg was then resealed and placed back into the 38°C incubator for 24h before collection. Ex Ovo electroporation followed a protocol similar to that developed by the Stern Lab (*88*). In short, HH9 embryos were isolated via Whatman filter paper rings, washed twice in Pannett Compton Saline (Table 3) and once in Tyrodes Saline (Table 3), and placed face down in an electroporation chamber with a plate electrode (made in house by the Crick Making Lab). Plasmid mixture was injected between the vitelline membrane and the embryo over the CLE. The second electrode was placed over the CLE of the embryo to achieve targeted plasmid electroporation (pulse regime - 7V, 50ms length, 3 pulses, 500ms interval). Embryos were then cultured for 24 hours using the EC culture method (*94*) in 30mm dishes at 38°C for 24 hours before collection.

**Table 3:**
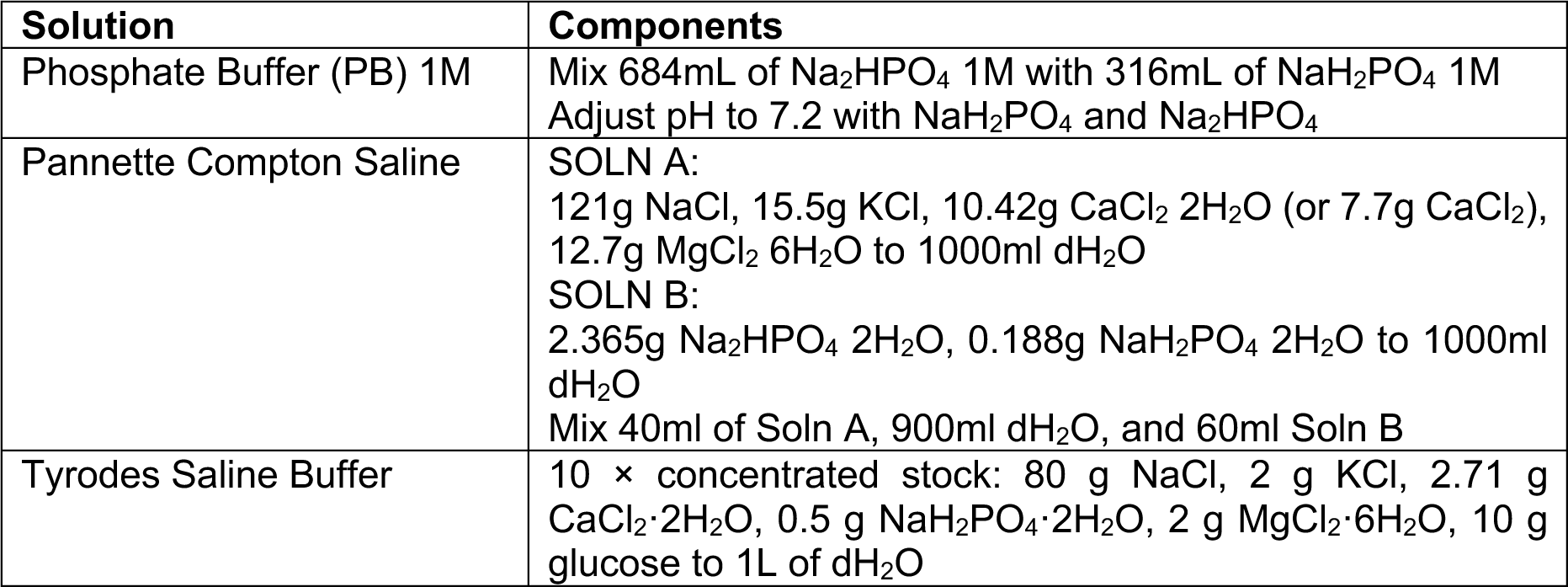
Solutions.

### Generation of CRISPR and RARA over-expression plasmids

The CRISPR system used was a modified version of that developed in the Sauka-Spengler Lab (*49*). Both the CAG-Cas9-2A-Citrine (Addgene: 92393) and pcU6_3-MS2-sgRNA (Addgene: 92394) plasmids were kindly donated by the Saurka-Spengler Lab, expanded, and maxi-prepped using a QIAGEN EndoFree Maxi Plasmid Kit. To modify the pcU6_3-MS2-sgRNA plasmid for capture sequencing, the 10X feature barcoding capture sequence 1 (Table 2) was inserted into hairpin 2 of the stem loop as described on the 10X platform website. To generate the multiple guide plasmids, the stock backbone plasmid was digested with Esp3I (ThermoFisher, ER0451), and double stranded oligos encoding each guide sequence were subsequently ligated into the vector, sequenced, and expanded as described in Williams et al.

RARα mutant over-expression plasmids (*80*) were kindly donated by Philippe Lefebvre and RARα sequences were PCR amplified with primers adding PaqCl cut sites (New England Biolabs, R0745S), cut via PaqCI and ligated into an EF1alpha-mScarlet backbone (*95*). Clones were subsequently sequenced, expanded, and purified via a QIAGEN EndoFree Maxi Plasmid Kit.

### Hybridization chain reaction (HCR)

All HCR followed the manufacturer protocol (Chicken embryos – Multiplexed HCR v3.0 protocol, Molecular Instruments). In short, HH10 embryos were fixed with 4% PFA for 1 hour and stored in Methanol overnight at −20°C. Embryos were then stepwise rehydrated by 25% steps of PBST (0.1% Tween 20, Sigma P2282) on ice, treated with protease K (VWR International, A3830.0500), and fixed again with 4% PFA for an additional 20 minutes at room temperature. Embryos were then transferred to 5x SSCT (SSC buffer with 0.1% Tween 20, Invitrogen 15557-044), pre-hybridized for 30min at 37°C with probe hybridization buffer (Molecular Instruments), incubated for 14 hours with 2pmol of probe, washed 4 times with 5x SSCT, pre-amplified with amplification buffer (Molecular Instruments) for 5 min at room temperature, incubated for 14 hours with hairpin solution in the dark at room temperature, and finally washed 4 times with 5x SSCT. Hairpins solution was prepared by snap cooling hairpins for 30mins and then mixing to generate a 60nM solution in amplification buffer. Embryos were then stained with DAPI for 20 minutes at room temperature, mounted on coverslips with ProLong™ Glass Antifade Mountant (Thermofisher, P36980), and subsequently imaged on a scanning confocal microscope (Leica SP8).

### HCR Quantification

HCR images were run through the analysis pipeline described in (*45*) to produce a matrix of cell centers, nuclear channel averages, hood channel averages, and cytoplasm channel averages. The hood averages which consist of a slight dilation of the nuclear mask were used as a proxy for total transcript detected per cell. Hood averages were then normalized to DAPI detection to account for imaging artifacts and cells with values larger than the mean of distribution of averages was determined to be positive signal in that channel (Figure S2B). Hierarchical clustering was performed on cell averages across the imaged channels to produce groups with overlapping HCR detection signatures (Figure 1G, S3A,B). From these groups the threshold of high TBXT (notochord) and medium TBXT (CLE/PS) was determined to be > 0.4 or >0.3 respectively. This was used to quantify the percent overlap of cells expressing varied levels of TBXT and each channel of interest.

### Embryo embedding, sectioning, and immunofluorescent staining

Embryos were either dissected from the eggs or from EC cultures, washed with PBS and fixed for 1 hour with 4% paraformaldehyde (16% diluted in PBS, Theromofisher Scientific, 28908). Embryos were then washed x3 in 0.24M phosphate buffer solution (PB)(Table 3) and left overnight in 0.12PB with 15% Sucrose (Sigma, 84100-1KG) at 4°C. Embryos were then embedded in gelatin (Sigma, G2500-500G), frozen in Isopentane (between −40°C and −50°C), and stored at −80°C. Blocks were sectioned on a Leica Cryostat CM3050S at 12um sections and placed on Superfrost Plus Adhesion Microscope slides (epredia, J1800AMNZ).

For immunostaining, slides were washed x3 with PBS at 42°C to remove gelatin, then blocked for 1 hour in PBS with 5% normal donkey serum (Merk Life Science, D9663) and 0.3% Triton^TM^ X-100 (Sigma-Aldrich, 1002135493), and incubated overnight with primary antibodies at 4°C (Table 4). Slides were then washed x4 with PBS, incubated with secondary antibodies (Table 4) for 1-2 hours at room temperature, incubated with DAPI for 15 minutes and washed with PBS. Finally, cover slides were mounted using ProLong™ Glass Antifade Mountant (Thermofisher, P36980), and subsequently imaged on a scanning confocal microscope (Leica SP8). For tortuosity measurements of neural tubes, images were processed in ImageJ to generate masks of the SOX2 HCR channel, marking the neural tube. Then with the Analyze Particles function, mask area and perimeter was determined for each neural tube.

**Table 4:**
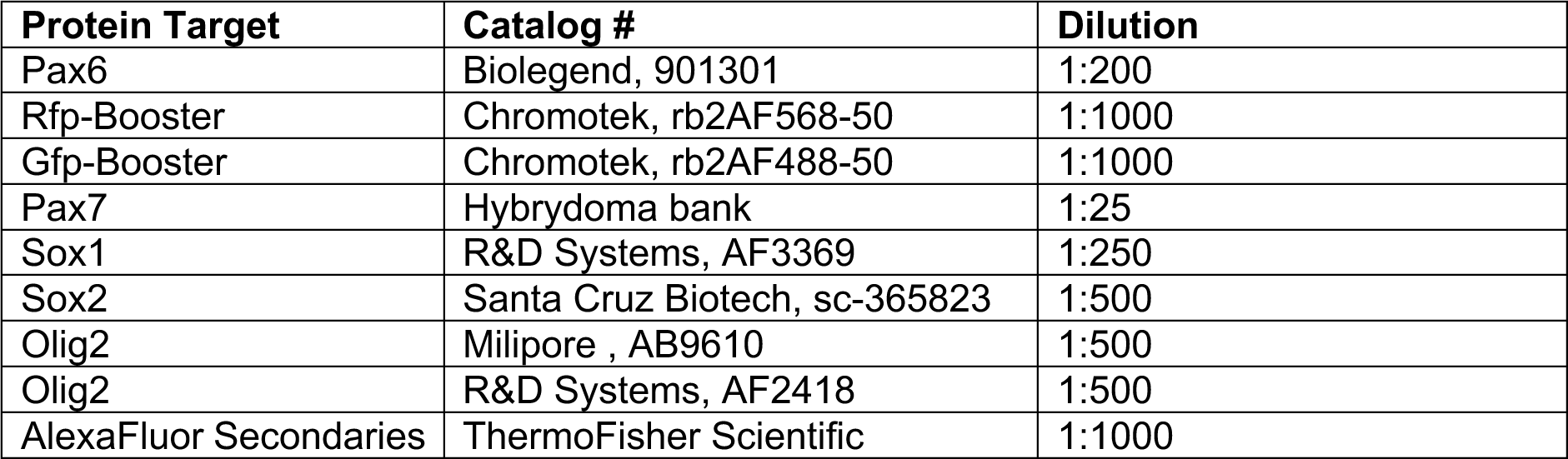
Antibodies.

### Flow Cytometry and Florescence Activated Sorting (FACS)

Embryos were isolated and dissociated to a single cell suspension by incubating in 200ul of dissociation solution consisting of Accutase (Stemcell Technologies, 07922) with 3U/mg Papain (Sigma-Aldrich, 10108014001) and 1mg/mL of Collagenase 4 (Gibco, 17104019) for 20 minutes at 38°C. Additionally, cells were briefly mechanically dissociated after the first 10 minutes incubation in dissociated solution with a P1000 pipet. Subsequently, 200μL of DPBS (Thermofisher Scientific, 14190144) was added, suspension filtered through a 40μm Flowmi cell strainer (Sigma-Aldrich, 136800040). Cells were spun for 4 minutes at 400g and resuspended in PBS+ 0.5% BSA (Sigma-Aldrich, A2153) and kept on ice for subsequent analysis.

For the cells intended for single cell sequencing experiments, Citrine+ cells were enriched via FACS using a BD Influx cell sorter with a 70um nozzle into DNA Lobind 1.5mL Eppendorf tubes (Merck, EP0030108051), spun post sort at 400g, and resuspended in 50ul of PBS+0.5% BSA for subsequent single cell transcriptomics (described transcriptomics methods section).

For fixed cell flow analysis, single cells dissociated and filtered were spun for 4 minutes at 400g and resuspended in 200μL DPBS with LIVE/DEAD^TM^ Fixable Dead Stain Near-IR fluorescent reactive dye (Themo Scientific, L34976), incubated on ice for 30 minutes in the dark, spun again for 4 minutes at 400g and finally resuspended in 100μL of 4% PFA for a 10 minute incubation at room temperature. Post incubation 1mL of DPBS was added, suspension spun at 2000g, resuspended in 500μL of DPBS + 0.5% bovine serum albumin BSA, and stored at 4°C.

For Flow analysis, cell suspensions were resuspended in PBS + 0.5%BSA and 0.1% Trition-X100 (X) with primary antibodies (Table 3) and incubated for 2h at room temperature, washed and then subsequently resuspended in a similar solution with secondary antibodies (Table 3) for 45min at room temperature. Finally, cells were washed once with PBS + 0.5% BSA and 0.1% Triton-X100 and resuspended in PBS+0.5% BSA and run on a BD LSRFortessa Cell Analyzer with subsequent analysis in FlowJo where population statistics were calculated via ANOVA and subsequent Tukey Tests.

### Single Cell RNA Sequencing Preparation

Single cell suspensions were counted post FACS and viability of >90% was ensured. A single cell suspension was loaded onto a Chromium Chip G and run on the 10x Chromium Controller (PN-1000120). The cells were then processed via the Chromium Next GEM Single Cell 3’ Kit v3.1, 16 rxns (PN-1000268) protocol and the 3’ Feature Barcode Kit, 16 rxns (PN-1000262). cDNA synthesis, library construction, and feature PCR amplification were performed via manufacturer’s protocol. Samples were then sequenced on an Illumina HiSeq4000 using 100 bp paired-end runs aiming for 50,000 reads per cell (between 30k-50k depending on sample).

### Data Processing of GEX and CRISPR Reads using Cell Ranger

All GEX datasets were processes using Cell Ranger (v4.0.0, 10X Genomics), where FASTQ files were generated via the cell ranger count pipeline and CRISPR barcodes were analysed via the CRISPR guide capture analysis pipeline. FASTQ were aligned to the white leghorn GRCg7w reference genome with the addition of the Citrine transgene. (https://www.10xgenomics.com/support/software/cell-ranger/latest/analysis/running-pipelines/cr-feature-bc-analysis)

For the reanalysis of the Rito et al. data set (accession number GSE223189), the 10 somite embryo data (GSM6940809) and the 13 somite embryo data (GSM6940810) were processed in the Cell Ranger count pipeline.

### Seurat Analysis of Rito et al. dataset

From the Cell Ranger pipelines, the Seurat package (version 5)(*96, 97*) was used in R (version 4.3.3) to analyse the resultant FASTQ files. For the Rito et al. dataset, removal of outliers for quality control included thresholds based on the numbers of genes/features (nCount_RNA > 2500, nFeature_RNA >300 & nFeature_RNA < 3500, mitochondrial gene percentage < 10% & mitochondrial gene percentage > 0.5%). Principal component analysis was used to identify the most variable genes and the top 30 principal components used to define a 16 cluster UMAP via the functions FindNeighbors, FindClusters, RunUMAP and DimPlot with a resolution boundary of 0.9. Using known marker genes (SOX2, TBXT, PAX6, MESP1, TBX6, NOTO, SOX10, FOXA2) the paraxial mesoderm, primitive streak, and neural tube lineages were identified and relevant clusters sub-setted into a new dataset using the subset function.

### Feature Selection Via Entropy Sorting

As an alternative to highly variable gene selection (HVG) for single cell RNA sequencing feature selection prior to downstream analysis, we used an Entropy Sorting (*46*) based feature selection algorithm, continuous Entropy Sort Feature Weighting (cESFW)(*47*). Using cESFW, we identified a set of 204 genes that facilitated the identification of 9 clusters of cells in the primitive streak, pre-neural tube and neural tube of the chick HH10 and HH13 single cell RNA sequencing data (Fig S1B). We then generated ranked gene lists for each of these 9 clusters using the Entropy Sort Score (ESS) correlation metric. The ESS metric is useful for identifying genes whose expression are specifically enriched in a population interest because it strongly penalizes positive gene expression that is found outside of the group of interest. The 9 identified clusters and ranked gene lists were found to be consistent with marker genes highlighted in the literature (Fig S1A,C). We then took the top 500 cESFW ranked genes in each cluster as candidates for possible genetic regulators of neural fate acquisition (Figure S1A,B). This identified 2505 differentially expressed genes (Table 1) as some genes overlapped between clusters. The workflow to reproduce these results may be found at the following Github repository, https://github.com/aradley/Libby-A-2024.

### Seurat Analysis of CRISPR Screen

Similar to the Rito et al. dataset, the screen single cell RNA sequencing dataset was first filtered based on the previously outlined thresholds for genes, features, and mitochondrial DNA percentage, we additionally filtered out a contaminating blood population via removing clusters that expressed high levels of the gene LMO2. Then the screen data was treated as a query dataset and mapped back to the wildtype dataset from Rito et al using the functions FindTransferAnchors, TransferData, AddMetaData, RunUMAP and MapQuery. This defined a UMAP transformation of the screen dataset to the original wildtype dataset. Cell cycle was scored using the function CellCycleScoring and subsequent scores were regressed to normalize the dataset. From here, we similarly subsetted the clusters associated with the paraxial mesoderm, primitive streak, and neural tube lineages and re-clustered following the previously described protocol using highly variable genes to define principal components and obtain a UMAP with 15 defined clusters. We subsequently excluded and LMO2+ (contaminating blood progenitors) leaving 14 total cell populations.

### Gene Enrichment Analysis

From the protospacer calls determined by the Cell Ranger CRISPR capture sequence pipeline, cell barcodes that had detected guides were identified and added to the meta data of the screen Seurat object. From this, the log10 UMI counts per cell of each protospacer were determined and two standard deviations from the mean log10 UMI count was discarded as ambient amplification. “Successful” guides were determined by using the function FindMarkers with the target CRISPR gene as the feature compared between the cells that had received the CRISPR gene of interest guide and those without. From this, 5-fold cross validation testing was used to generate enrichment scores of each individual guide’s representations per cluster where guide representation was calculated as total cells with guide in cluster divided by the total cells within the cluster. Chi-squared tests comparing the average enrichment across all guides per cluster, scramble 1 enrichment, and the target gene of interest enrichment were used to determine which guides’ enrichment patterns deviated from the scramble guide (z-score >2 and <-2). This was confirmed by Kullback-Leibler Divergence cross validation tests using the Philentropy R package (p< 0.05)(*98*). K-means clustering of the enrichment patterns was determined by minimizing the intra-cluster distances via elbow plot.

### Differential Gene Expression and GO Term Analysis of Screen

To determine the effect of each gene perturbation beyond reduced expression of the targeted gene the scMAGeCK-LR package (*67*) was used to determine differentially expressed genes post perturbation compared to the scramble control. Here the 2502 genes from Entropy Sort Score ranked gene list were used as the features of the input Seurat object and only genes with a p<0.05 were considered differentially regulated post perturbation. The Enrichr package (*99–101*) was then used to identify significant Gene Ontology terms (“GO_Biological_Process_2015”) and KEGG Pathways (KEGG_2021_Human) with adjusted p-values < 0.05.

### Data and Code availability

Single-cell RNA sequencing data have been deposited onto Gene Expression Omnibus under the ascension number GSE267653. The transcriptomics analysis scripts are available in Git hub repositories: https://github.com/aradley/Libby-A-2024 and https://github.com/Libbya-crick/Libby-2024-CRISPy-Chickens. Generated plasmids have been deposited on Addgene under the following codes: 92393 and 84348.

---

**Figure S1: Establishment of dataset genes of interest and image analysis**

A) Heat map showing top 10 ranked Entropy Sort Scored genes across the wildtype dataset where rows refer to pseudo-bulk regions of the reference UMAP in Figure 1B (PS – primitive streak, PreNT – preneural tube, NT – neural tube, NC – neural crest, FP – floor plate). B) UMAP showing cESFW driven clustering of wildtype dataset that was used to define the top 500 genes per cluster. (PS – primitive streak, PreNT – preneural tube, NT – neural tube, NC – neural crest, FP – floor plate) C) Pseudo-lineage trajectories used to generate the heatmap used in Figure 1E. Two trajectories depicted ending in either the neural crest population or the floor plate population. D) UMAPs depicting the gene expression of known lineage marker genes across the wildtype dataset clustered using Louvain clustering based on top ranked Entropy Sorted genes. Bottom right UMAP shows neuromesodermal progenitors identified by the co-expression of *SOX3/2* and *TBXT* expressing cells.

**Figure S2: Quantification of Hybridized Chain Reaction (HCR) images**

A) Example image analysis segmentation for quantification of HCR images. B) Example thresholding based on mean values of each channel. Red line indicates the mean. Grey lines indicate one standard deviation away from the mean and 5 standard deviations away from mean. C) Quantification maps from the xy plane of *TBXT, SOX2,* and *F2RL1* HCR in HH10 embryos where gene expression detection is normalized to DAPI detection (rostral top-caudal bottom). D) Quantification maps from the xy plane of *TBXT, SOX21,* and *MLLT3* HCR in HH10 embryos where gene expression detection is normalized to DAPI detection in the xy plane (rostral top-caudal bottom). E) Quantification maps from the xy plane of *TBXT* and *CMTM8* HCR in HH10 embryos where gene expression detection is normalized to DAPI detection (rostral top-caudal bottom).

**Figure S3: Population mapping from HCR quantification to embryo and single cell dataset**

A) Hierarchical clustering of *TBXT* transcript count with that of *SOX21, MLLT3, CMTM8*, and *F2RL1*. Light blue dots highlight population plotted in Figure 1I or S3B. B) Mapping of *SOX21*+ light blue populations from Figure S3A (blue dots) onto *TBXT* expression maps of the imaged embryo in the xy plane (rostral top-caudal bottom). C) Mapping of *SOX21*+ or *CMTM8+* populations in light blue from Figure S3A onto *TBXT* expression maps of the imaged embryo in the yz plane (rostral-left caudal-right, NT – neural tube, PS – primitive streak, NOTO - notochord). D) Mapping of *MLLT3*+ or *F2RL1+* populations in light blue from Figure S3A onto *TBXT* expression maps of the imaged embryo in the yz plane (rostral-left caudal-right, NT – neural tube, PS – primitive streak, NOTO - notochord). E) Quantification of cells co-expressing query genes with low, medium, and high levels of *TBXT* expression in quantified HCR images. I) UMAPs of the overlapping expression domains of *MLLT3, F2RL1,* and *CMTM8* with reported markers of CLE and neural tube domains *ADAMTS19, CNTN2, NEFM*.

**Figure S4: Flow cytometry of OLIG2 knockout embryos**

A) Immunofluorescent images for the indicated makers in transverse sections of the neural tube 24h post electroporation. White arrows mark detection of Cas9 and in KO condition loss of Olig2 protein. B) Example flow gates for quantification of Sox2+ progenitors in embryos and subsequent thresholding for Olig2+ cells.

**Figure S5: CRISPR screen dataset analysis and quantification**

A) UMAPs depicting the gene expression of known lineage marker genes across the screen RNA sequencing dataset. B) Quantification of the distribution of detected guide UMI counts per cell. Red line marks 2 standard deviations from the mean where guide detection below is considered random RNA amplification.

**Figure S6: Analysis of screen guide enrichment and gene expression**

A) Quantification of Kullback-Leibler divergence tests of each knockout compared to the distribution of Scramble guide 1. (*,**,*** indicates p = 0.05, p = 0.001, and p = 0.0001, respectively by ANOVA and subsequent Tukey tests) B) Heatmap showing calculated Jaccard distances between gene perturbations based on KEGG pathways called by each knockouts differentially expressed gene list. B) Elbow plot of within cluster sum of square (wcss) as the number of k-means clusters increases when applied to differential gene expression across knockouts.

**Figure S7: MLLT3, CMTM8, GREB1, and SPRY2 perturbed cells show overlapping differentially regulated genes**

A) Heatmap displaying the overlapping differentially regulated genes between *MLLT3, CMTM8, GREB1*, and *SPRY2* perturbed cells (grey = no differential expression by scMAGeCK algorithm, i.e. p>0.05 by FDR).

**Figure S7: Quantifications of dual mScarlet and Citrine electroporated embryos**

A) Z-projection of whole mount fluorescence of scramble and *MLLT3* knockout embryos using HCR for *SOX2, TBXT, MLLT3*, and reporter fluorescence of Citrine (marking Cas9 and presumed KO). *(top)* dorsal view of embryo; *(bottom)* ventral view of embryo. B) Z-projection of whole mount fluorescence images of scramble and *MLLT3* knockout embryos using HCR for *PAX6* and *MLLT3*. C) Example flow plot of SOX2+ progenitors of either scramble control (grey) or *MLLT3* knockout (pink) where quadrants indicate presence of Citrine or mScarlet populations. D) Quantification of flow cytometry of dually electroporated embryo knockouts showing ratio of Citrine to mScarlet across all cells indicating no significant loss of Citrine cells that have received a targeting guide compared to non-targeting. (n.s. represents non-significance by ANOVA, n = 4-7 embryos per condition).

**Sup. Table 1: List of Entropy Sort Score ranked genes**

Provided Excel file

