## Supplementary figures and images for "An in vivo CRISPR screen in chick embryos reveals a role for MLLT3 in specification of neural cells from the caudal epiblast"

### Supp. Figures

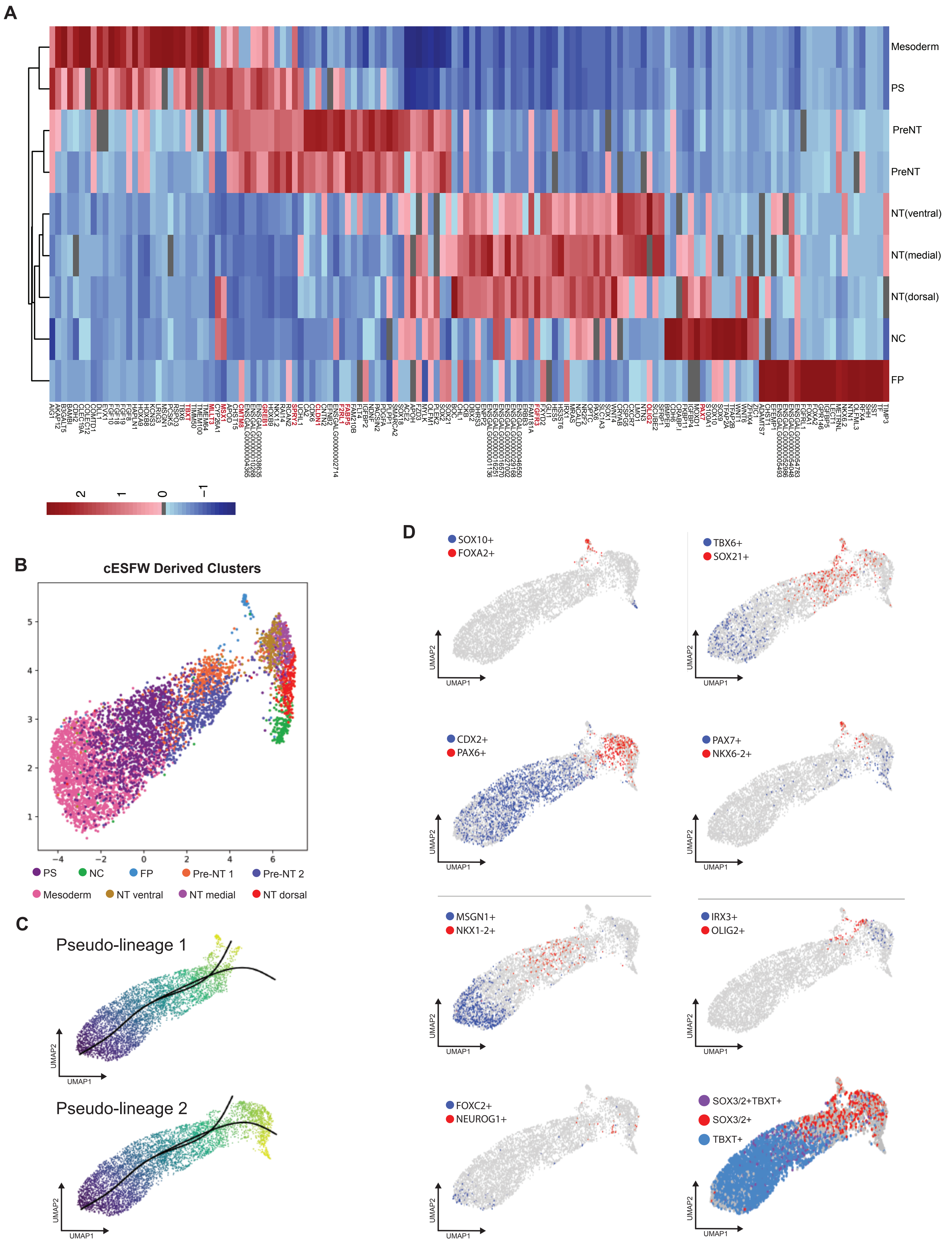

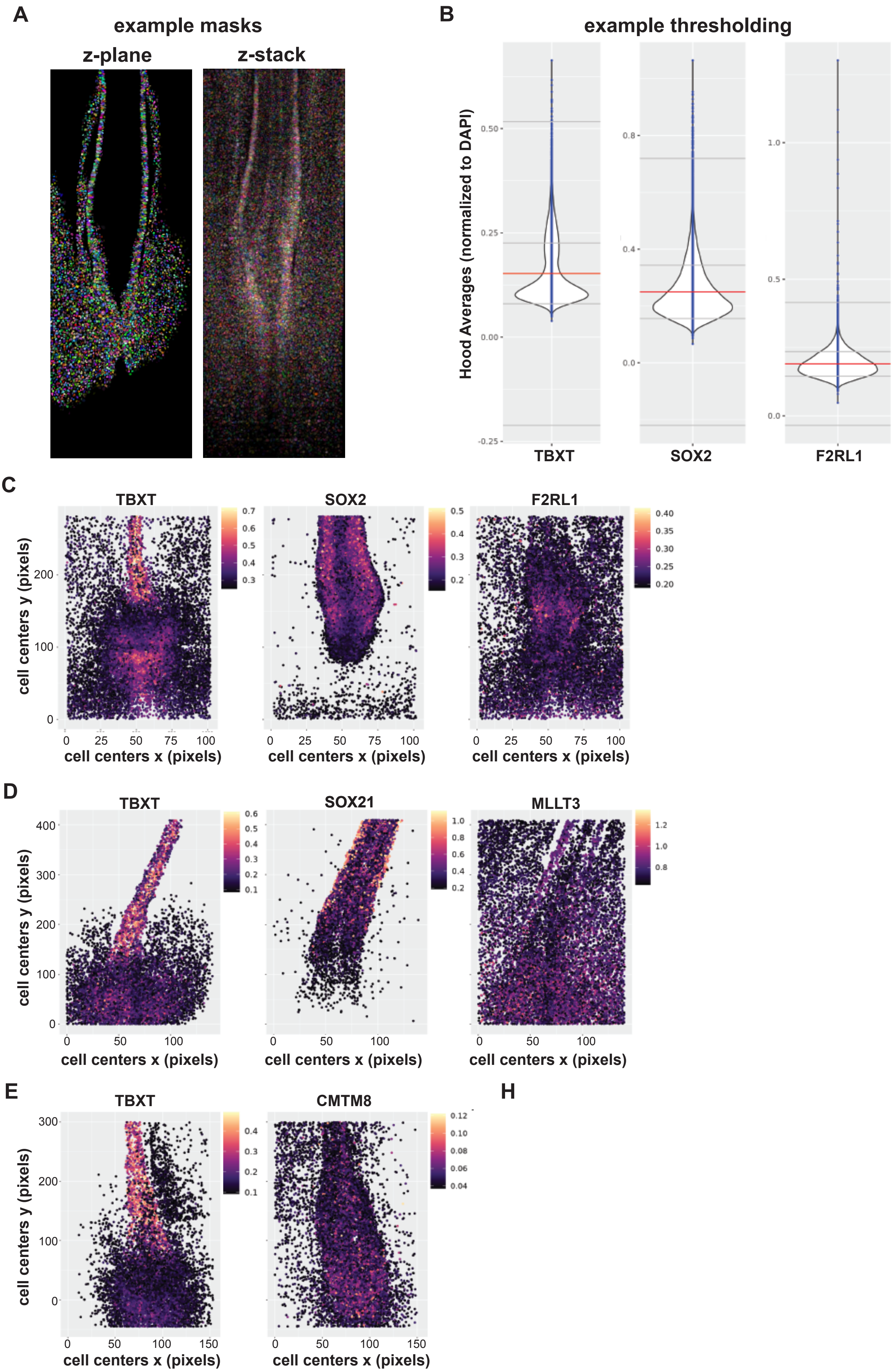

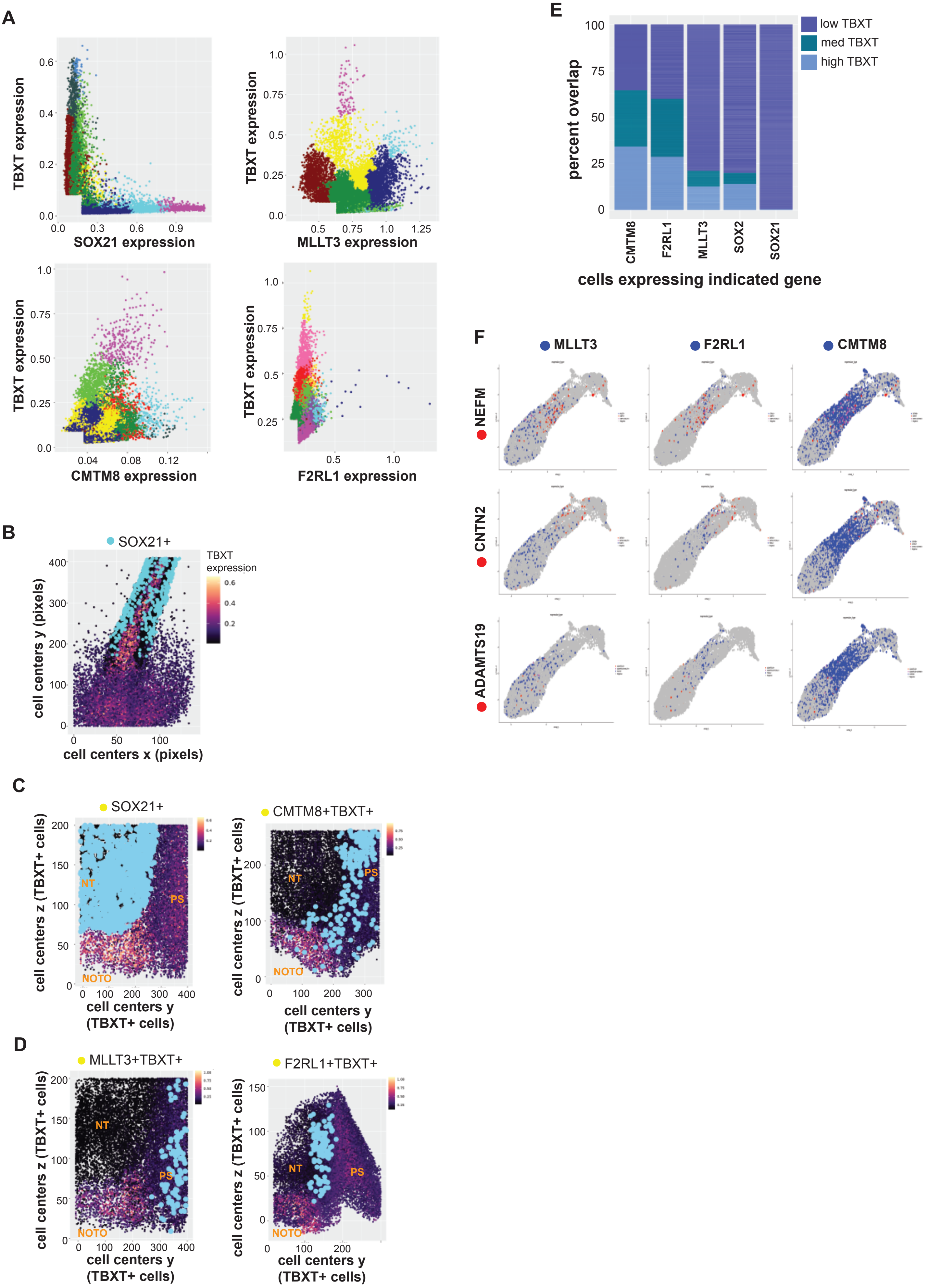

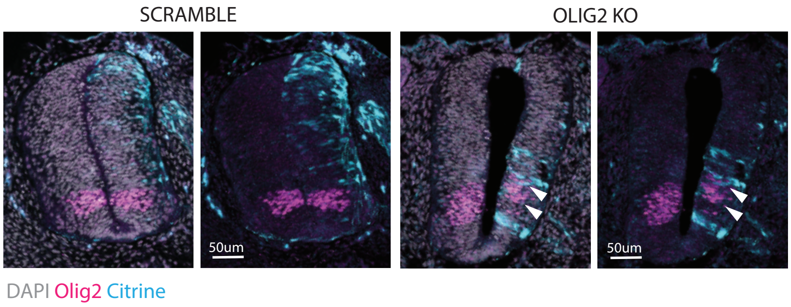

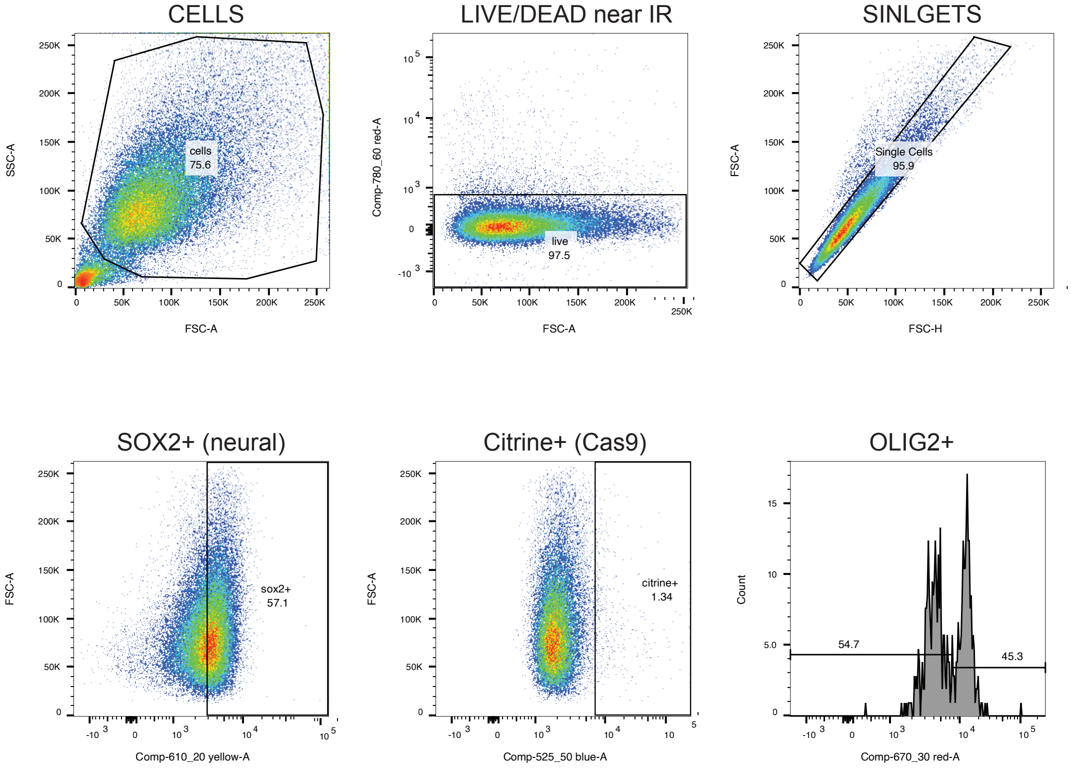

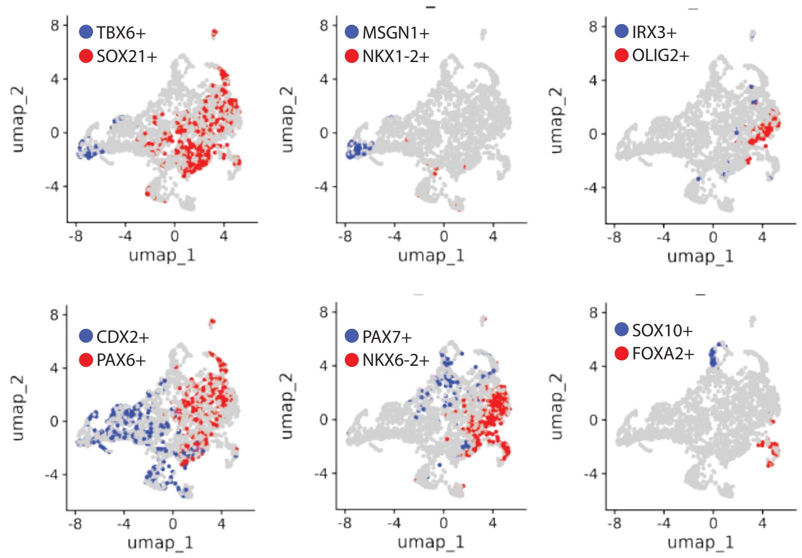

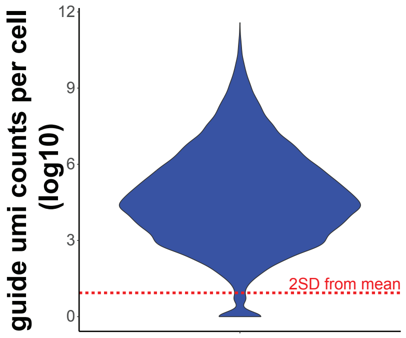

#### FIGURE S6

**A**

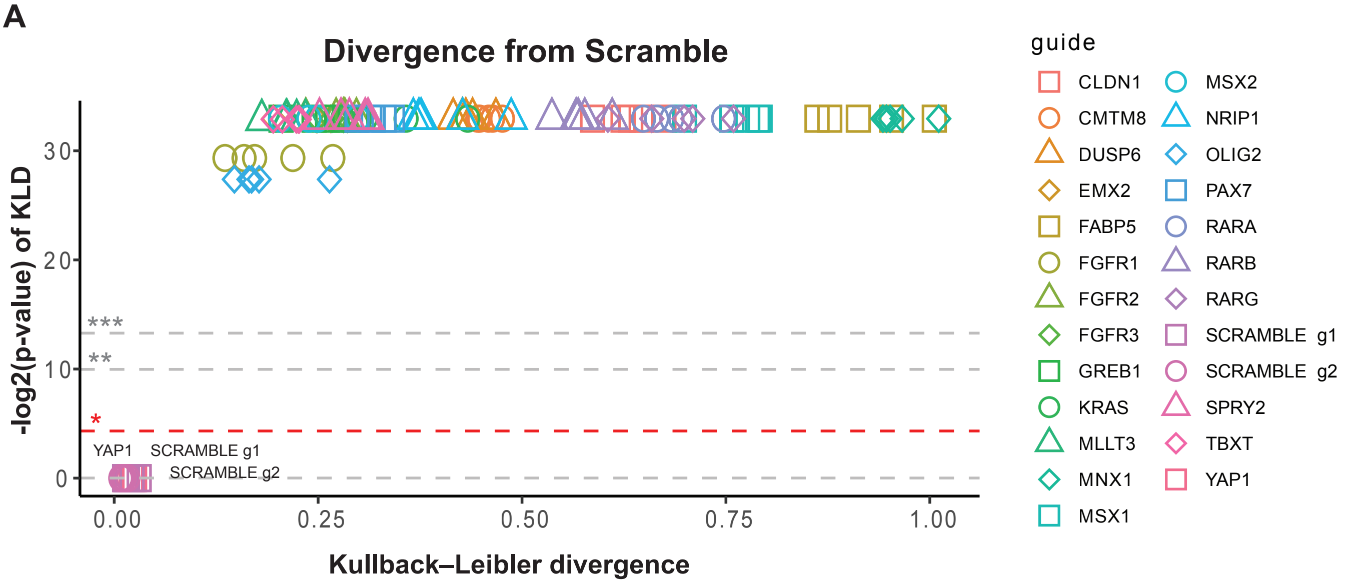


**B**

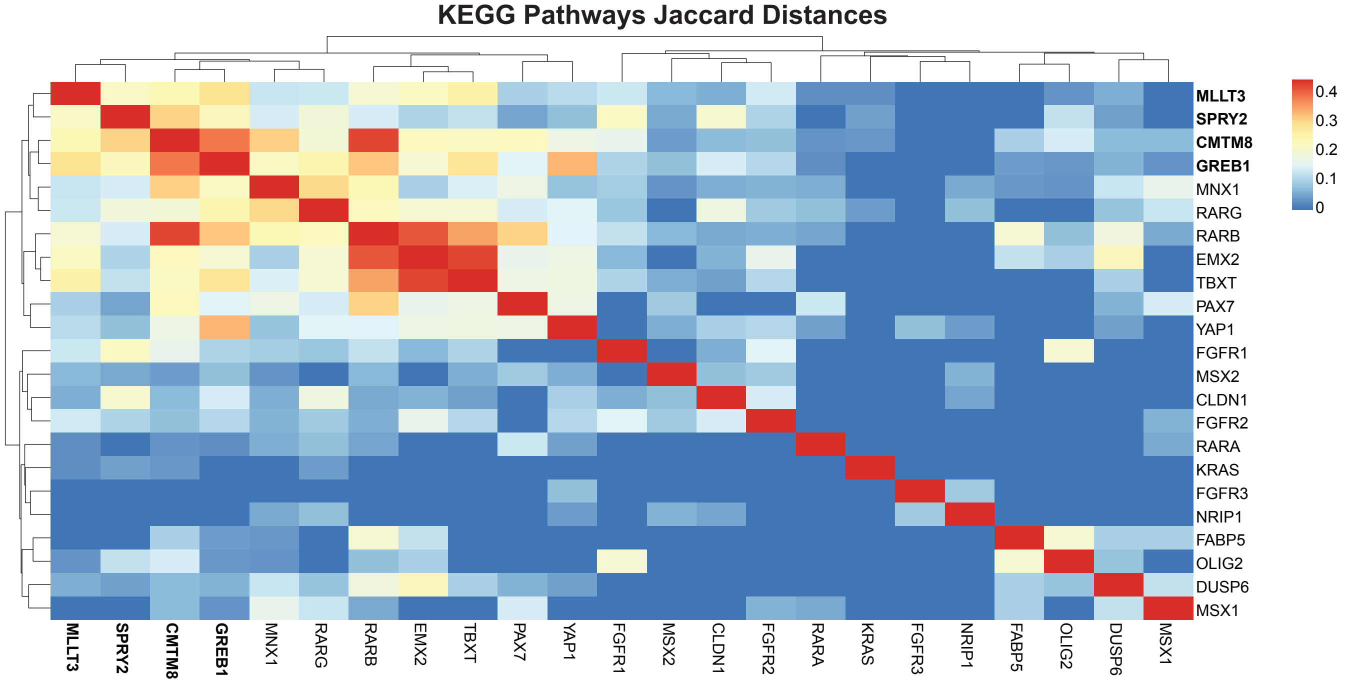


**C**

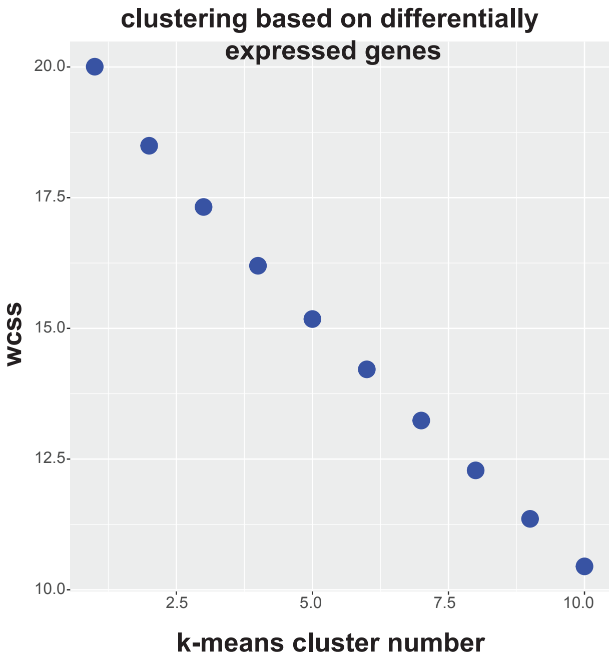

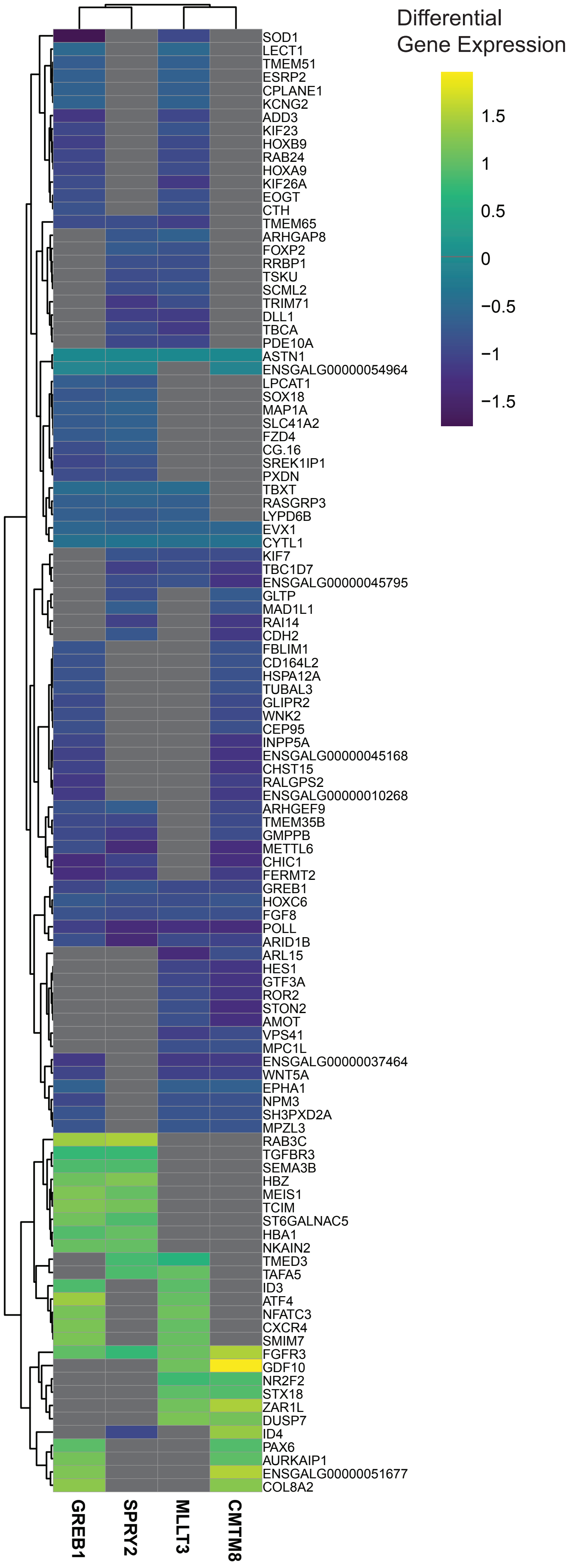

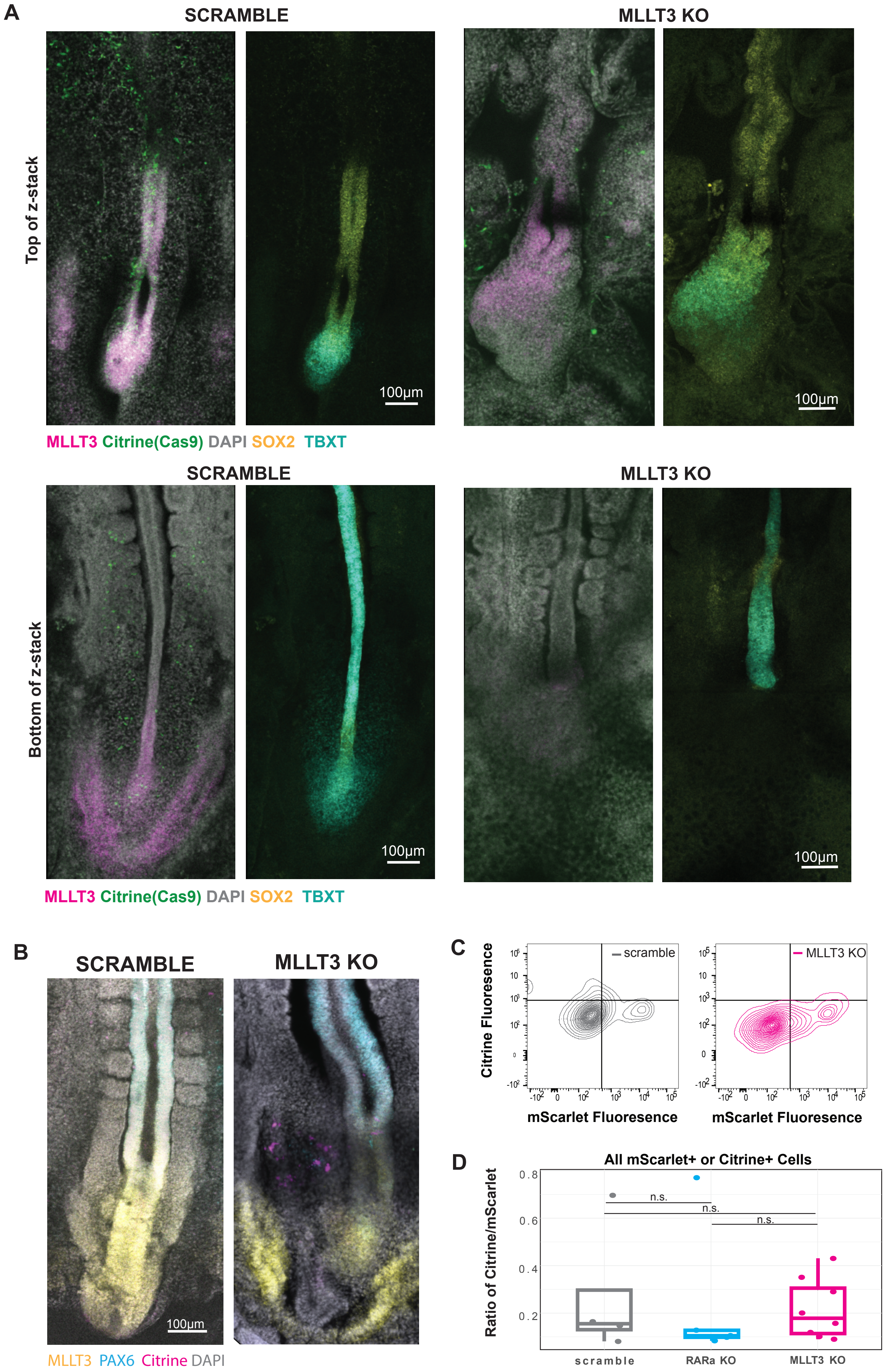
